# Nav1.2 and BK channels interaction shapes the action potential in the axon initial segment

**DOI:** 10.1101/2022.04.12.488116

**Authors:** Luiza Filipis, Laila Ananda Blömer, Jérôme Montnach, Michel De Waard, Marco Canepari

**Author notes:** **Address of the submitting and corresponding author:** Marco Canepari, Laboratoire Interdisciplinaire de Physique (UMR 5588), Bat. E45, 140 avenue de la physique, Domaine univ., 38402 St Martin d’Hères cedex, France. These authors contributed equally (Luiza Filipis, Laila Ananda Blömer). **Author contributions:** L.F., L.A.B. and M.C. designed the research, L.F. and L.A.B. performed brain slice research and data analysis, J.M. and M.d.W. designed the novel peptide and characterised ion channel selectivity of peptides, L.F. developed the NEURON model and performed computer simulations, M.C. wrote the paper. All authors have given approval to the final version of the manuscript. **Competing interests:** The authors declare no competing interests.

## Abstract

In neocortical layer-5 pyramidal neurons, the action potential (AP) is generated in the axon initial segment (AIS) when the membrane potential (V_m_) reaches the threshold for activation of two diverse voltage-gated Na^+^ channels, that differ in spatial distribution and biophysical properties. Yet, the understanding of specific functional differences between these two channels remains elusive. Here, using ultrafast Na^+^, V_m_ and Ca^2+^ imaging in combination with partial block of Na_v_1.2 by a recent peptide, we demonstrate an exclusive role of Na_v_1.2 in shaping the generating AP. Precisely, Ca^2+^ influx *via* Na_v_1.2 activates BK Ca^2+^-activated K^+^ channels determining the amplitude and the early phase of repolarisation of the AP in the AIS. We mimicked this result with a NEURON model where the role of the different ion channels tested reproduced the experimental evidence. The exclusive role of Na_v_1.2 reported here is important for understanding the physiology and pathology of neuronal excitability.

## Introduction

In most mammalian neurons, the action potential (AP) is generated in the axon initial segment (AIS) where its waveform is shaped [1]. After generation, the AP propagates along the axonal branches, regenerates at Ranvier nodes, and the waveform at synaptic terminals regulates neurotransmitter release providing an analog modulation of synaptic transmission [2]. The AP from the AIS also back-propagates to the soma and then along the dendrites, progressively changing its shape and regulating synaptic integration [3]. Eventually, the AP waveform in the AIS sets the way in which a neuron can respond to prolonged depolarisation by firing several APs at high frequencies, determining the ability of the neuron to encode signals [4]. The build-up of the AP waveform in the AIS is the result of the sequential activation, inactivation and de-activation of diverse voltage-gated ion channels [5]. In this cascade of events, the activation of voltage-gated Na^+^ channels (VGNCs) is the first step. In the AIS of excitatory pyramidal neurons, Na_v_1.2 and Na_v_1.6 are the two isoforms of VGNCs present [6] and the understanding of the functional role of these two channels is therefore a fundamental piece of information necessary to link the AP shape to the later sequence of ion-channel activation and de-activation steps. Specifically, in neocortical layer-5 (L5) pyramidal neurons, these two channels have different axonal distributions [7] and biophysical properties [8]. The Na^+^ current associated with the AP is characterised by a fast inactivating component and by a non-inactivating component [9]. Na_v_1.6-deficient mice are characterized by a reduced non-inactivating Na^+^ current [10], whereas Na_v_1.2-deficient mice are characterized by reduced dendritic excitability and synaptic function [11]. Finally, it was recently found that Na_v_1.2 channels are not only permeable to Na^+^, but also to Ca^2+^, and that a Ca^2+^ influx mediated by VGNCs occurs during the AP in the AIS [12]. The precise native role of Na_v_1.2 and Na_v_1.6, however, and their contributions to AP generation remains to be understood. Dysfunctions of these channels are the cause of neuronal disorders in patients carrying critical mutations in their encoding genes. In the case of Na_v_1.2, several gene variants cause diverse neuropsychiatric syndromes including epilepsies with different degree of severity, intellectual disability and autism with or without epileptic seizures [13]. Also, channelopathies of Na_v_1.6 are associated with rare but severe cases of epileptic encephalopathy [14]. Clearly, shedding light on the signaling processes triggered by VGNCs is an important step for the development of innovative therapeutic strategies to tackle these genetic pathologies, but this goal requires first a thorough understanding of the contributive role of each channel subtype to neuronal excitability.

Until recently, the study of the activation and signaling of one specific VGNC isoform has been limited by a series of experimental obstacles. First, although partially selective, VGNC blockers act on several isoforms and acceptable discriminative selectivity between Na_v_1.2 and Na_v_1.6 could not be achieved until now. Second, the inhomogeneous distribution of VGNCs, as well as the site-dependence of the AP waveform, requires recording ion concentrations and V_m_ transients at high spatial and temporal resolution for a proper understanding of the role of each channel subtype. Finally, the change of membrane potential (V_m_) transients induced by VGNC inhibition affects the activation of the other voltage-gated ion channels involved in the AP shaping. This important synergy of diverse ion channels, underlying the AP generation, makes the combined analysis of all these channels necessary. In this study, we tackled all these challenges and analysed the role of native Na_v_1.2 in the generation of the AP. This analysis was grounded on a partial but selective and stable effect on Na_v_1.2, with respect to Na_v_1.6, using a recent peptide mutated from a wild-type animal toxin. In the AIS of L5 pyramidal neurons, we optically measured Na^+^ currents [15], V_m_ transients [16] and Ca^2+^ currents [17-19] associated with APs elicited by somatic current injections and we analysed how the partial block of Na_v_1.2 affected these signals. We then focussed on the changes of these signals produced by fully blocking voltage-gated Ca^2+^ channels (VGCCs) and/or Ca^2+^-activated K^+^ channels (CAKCs) in order to correlate Na_v_1.2 signalling with the activation of these channels. To assess whether the role of Na_v_1.2 was exclusive, we performed control experiments using the partially selective Na_v_1.6 inhibitor 4,9-anhydrotetrodotoxin. Finally, in order to reconstruct the sequence of activation of the channels underlying the shaping of the generating AP, we built a NEURON model starting from the experience of existing models of the AIS of L5 pyramidal neurons [7, 12, 20-24], available in the NeuroDB database. The strategy, already used to realistically reconstruct the activation of the diverse dendritic ionic currents following a climbing fibre synaptic potential in cerebellar Purkinje neurons [25], consisted in reproducing the experimental profiles of ionic and V_m_ transients and their modifications following selective blocks of individual channel types. In this way, we could mimic the experimental effect of partially blocking Na_v_1.2 on the shape of the AP in the AIS.

## Results

### G^1^G^4^-huwentoxin-IV is selective for Na_v_1.2 against Na_v_1.6

The first necessary condition to unambiguously investigate the role of Na_v_1.2 in the AP generation at the AIS of L5 pyramidal neurons is the availability of a selective blocker of this channel that is specifically inert with respect to Na_v_1.6. Phrixotoxin-3 [26] has been reported to be selective for Na_v_1.2, but when we tested this peptide in L5 pyramidal neurons we found variable and rapidly reversible effects at the lowest concentrations where this selectivity is expected to apply. Thus, to produce a less reversible and still selective effect on Na_v_1.2 channels, we used a peptide modified from huwentoxin-IV [27] where the two glutamate residues at positions 1 and 4 were replaced with two glycine residues. The peptide (G^1^G^4^-huwentoxin-IV or simply hwtx herein) exhibits higher selectivity for Na_v_1.2 with respect to Na_v_1.6 at nanomolar concentrations as measured in HEK293 cells using automated patch-clamp recordings (Supplementary Fig. S1). Specifically, at 1 nM and 3.3 nM, hwtx blocks 16 ± 7 % and 53 ± 7 % Na_v_1.2 currents respectively, and only 5 ± 4 % and 9 ± 5 % Na_v_1.6 currents respectively. The challenge remained to exploit this selectivity by locally blocking Na_v_1.2 in the AIS of L5 pyramidal neurons in the context of brain slices in which effective concentrations may differ. In this study, we used the experimental configuration illustrated in Supplementary Fig. S2a. L5 pyramidal neurons were patched in the cell body and another pipette near the AIS was used to locally deliver selective channel blockers, for 1 minute, by gentle pressure application. In this way, when we delivered hwtx at 40 nM, the peptide did not affect the AP (Supplementary Fig. S1). The smallest concentration at which we consistently observed an affect was 80 nM and this effect was a delay in the AP onset and peak with occasionally a small decrease in amplitude (Supplementary Fig. S1). Finally, at 160 nM, hwtx consistently produced a longer delay and decreased the amplitude of the AP. Importantly for this project, at all concentrations, the consistent effects of hwtx were stable for several minutes after delivery, in sharp contrast to the case of phrixotoxin application. We could therefore use hwtx at 80 nM concentration to investigate Na^+^, V_m_ and Ca^2+^ signals associated with an AP, on the hypothesis that at this minimal concentration the peptide should produce a partial but selective block on Na_v_1.2 because this channel type is more sensitive than Na_v_1.6 to hwtx block. To this purpose, the protocol that we used in this study for ultrafast imaging from the AIS at 10 kHz is illustrated in Supplementary Fig. S2b. Given the morphological variability, we standardised our analyses to regions of 5 μm lengths within the proximal (prox) part of the AIS, at 5-15 μm from soma, indicated with “*1*” in all figures, and to regions of 5 μm lengths within distal (dist) parts of the AIS, at 30-40 μm from soma, indicated with “*2*” in all the figures. Fig. 1a shows the AIS of a neuron filled with the Na^+^ indicator Ion Natrium Green-2 (ING-2).

**Fig. 1.**
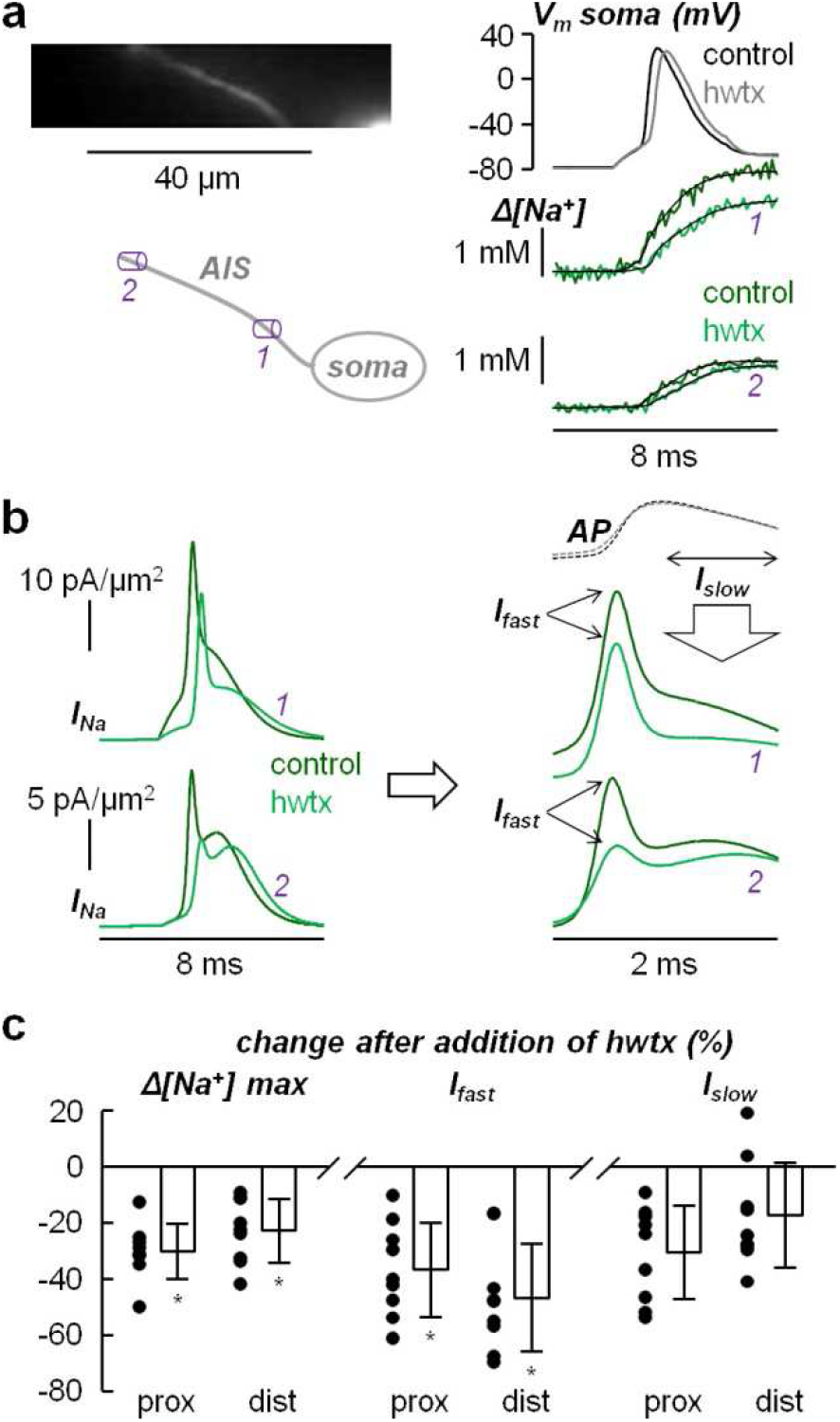
Effect of hwtx (block of Nav1.2) on the Na^+^ influx in the AIS. **a**, Left, fluorescence image (Na^+^ indicator ING-2) of the AIS and its reconstruction with a proximal region (*1*) and a distal region (*2*) indicated. Right, Somatic AP in control solution and after local delivering 80 nM of hwtx (top) and associated corrected Na^+^ transients fitted with a model function in *1* and *2*. **b**, Left, from the experiment in a, Na^+^ currents calculated from the time-derivative of the Na^+^ transient fits. Right, zoom of the Na^+^ currents 1 ms before and 1 ms after the AP peak indicating the fast component of the Na^+^ current (I_fast_) and the slow component of the Na^+^ current (I_slow_) defined as the mean Na^+^ current during the 1-ms interval after the AP peak. **c**, Left, single cell values and percentage change (N=9 cells) of the Na^+^ transient maximum after locally delivering of hwtx in proximal regions (mean ± SD = -30.3 ± 9.8), and in distal regions (mean ± SD = -22.9 ± 11.4). Centre, single cell values and percentage change of I_fast_ after locally delivering hwtx in proximal regions (mean ± SD = -36.6 ± 16.7), and in distal regions (mean ± SD = -46.8 ± 19.2). Right, single cell values and percentage change of I_slow_ after addition of hwtx in proximal regions (mean ± SD = -30.7 ± 16.6), and in distal regions (mean ± SD = -17.5 ± 18.6). “*” indicates a significant change (p<0.01, paired t-test).

The sodium concentration change (Δ[Na^+^]) associated with an AP was quantitatively measured as previously described [15] and the Na^+^ current (I_Na_) was calculated as the time-derivative of the fit of the Δ[Na^+^] signal using a model function (Fig. 1b). In the example of Fig. 1a, application of 80 nM hwtx produced a decrease of ∼30% of the Δ[Na^+^] signal in the proximal part of the AIS and a ∼15% decrease in the distal part. As already reported [15], the I_Na_ is characterised by a fast inactivating component (I_fast_) and a slow non-inactivating component (I_slow_). Thus, we systematically evaluated the change in the I_fast_ peak and in the I_slow_ (defined as the mean I_Na_ in the millisecond following the AP peak) after hwtx. On average, in 9 cells analysed, we found a significant (p<0.01, paired t-test) decrease in the Δ[Na^+^] and in the I_fast_ signals, both in proximal and distal regions, whereas a consistent but more variable decrease in the I_slow_ signals was estimated. In proximal regions, where Na_v_1.2 is expected to be dominant, the decrease in Δ[Na^+^] and I_fast_ signals were 30% and 36% respectively, indicating a block of 30-40% of Na^+^ channels. According to the selectivity curve reported in Supplementary Fig. S1, this percentage block is within the range where hwtx is inert to Na_v_1.6.

### The selective block of Na_v_1.2 widens the generating AP waveform in the AIS

The important finding that local delivery of 80 nM hwtx blocks 30-40% of Na_v_1.2 without affecting Na_v_1.6 allowed us performing further experiments with high confidence using hwtx at 80 nM. It permitted the analysis of the axonal AP waveform change after reducing Na_v_1.2, using V_m_ imaging with the voltage-sensitive dye JPW1114. In the example of Fig. 2a, application of 80 nM hwtx delayed the onset of the AP, consistently to what we reported in Fig. 1 and in Supplementary Fig. S1.

**Fig. 2.**
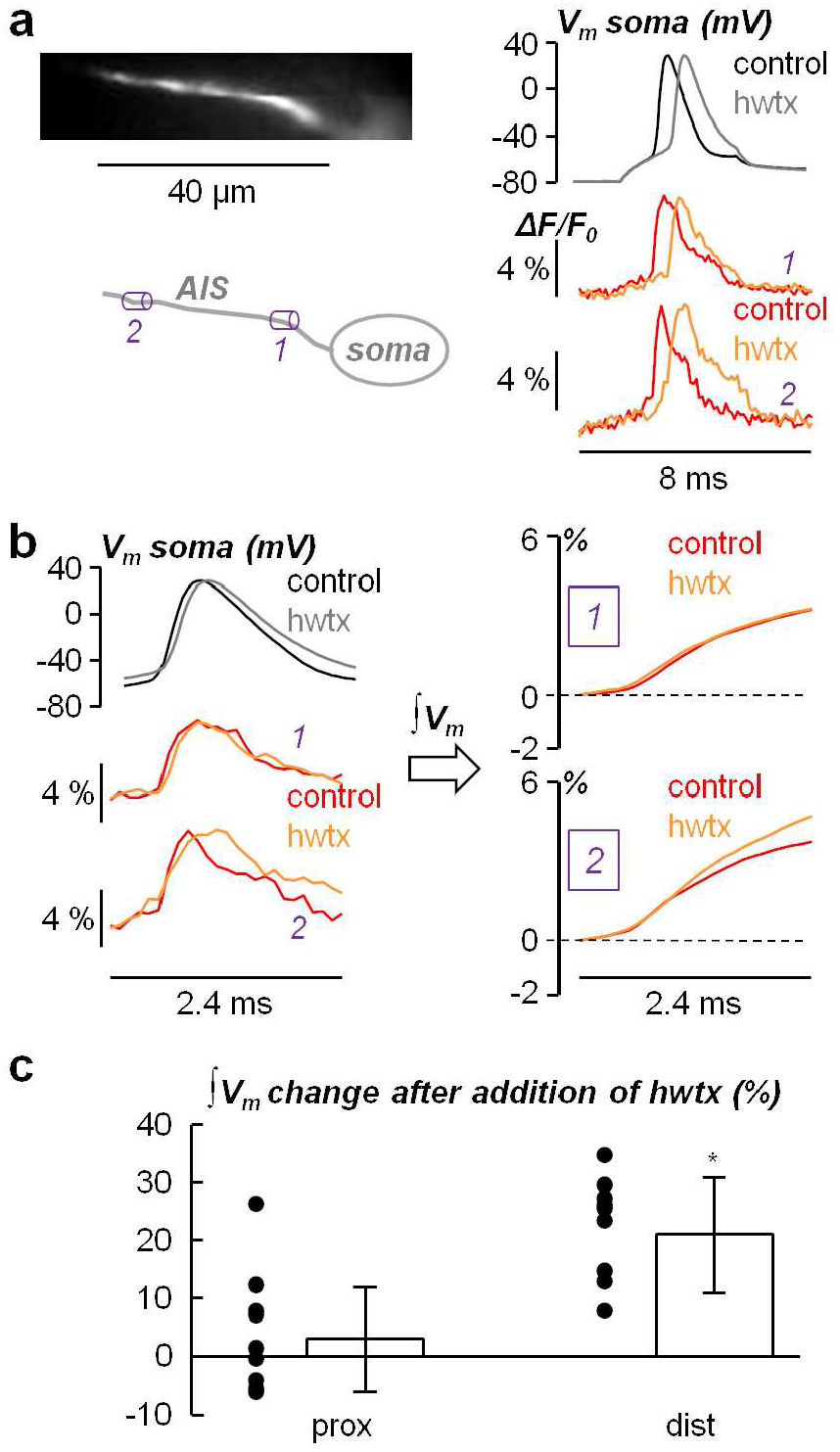
Effect of hwtx (block of Nav1.2) on the AP waveform in the AIS. **a**, Left, fluorescence image (voltage sensitive dye JPW1114) of the AIS and its reconstruction with a proximal region *(1)* and a distal region (*2*) indicated. Right, Somatic AP in control solution and after locally delivering 80 nM of hwtx (top) and associated V_m_ transients in *1* and *2*. **b**, Left, somatic and axonal APs at a different time scale (2.4 ms duration) visually indicating a widening of the AP waveform, in the distal position only, after locally delivering hwtx. Right, quantification of the AP waveform shape by calculation of V_m_ integral (∫V_m_) over the 2.4 ms time window comprising the AP signal. **c**, Single cell values and percentage change (N=9 cells) of the ∫V_m_ signal maximum after locally delivering hwtx in proximal regions (mean ± SD = 3.2 ± 8.8), and in distal regions (mean ± SD = 21.3 ± 10.5). “*” indicates a significant change (p<0.01, paired t-test).

In the distal AIS, however, it also changed the shape of the AP by widening its waveform (Fig. 2b). To reliably quantify the waveform change beyond the noise limitations, we calculated the time integral of the V_m_ signal in control conditions and after hwtx application over a time-window of 2.4 milliseconds containing the AP (∫V_m_). Notably, in 9 cells analysed, the ∫V_m_ at its last value significantly increased by 21.3 ± 10.5 % (p<0.01, paired t-test) in the distal regions of the AIS (Fig. 2c). This result indicates that Na_v_1.2 is contributing to the shaping of the generating AP by sharpening its waveform. The widening of the AP waveform, however, cannot be produced by a reduction of a Na^+^ (inward) current. The phenomenon reported in Fig. 2 can only be explained if the partial block of Na_v_1.2 is also associated with a reduction of the outward current which must be larger than the Na^+^ current change in the early phase of the AP repolarisation. The output current underlying the AP repolarisation is mediated by K^+^ channels. Since the AP amplitude is not significantly affected by the partial block of Na_v_1.2, any variation of current mediated by voltage-gated K^+^ channels (VGKCs) is minimal. The most likely hypothesis is therefore that CAKCs are also contributing to the shaping of the generating AP and that these channels are activated by Ca^2+^ influx *via* Na_v_1.2. Since Na_v_1.2 is also permeable to Ca^2+^ and it was reported that Na_v_1.2 mediates Ca^2+^ influx in the AIS during APs [12], we next investigated this phenomenon in detail using hwtx.

### Na_v_1.2 mediates Ca^2+^ influx associated with the AP in the AIS

To investigate Ca^2+^ influx and currents associated with an AP in the AIS, we recorded Ca^2+^ fluorescence from the indicator Oregon Green BAPTA-5N (OG5N) and estimated the Ca^2+^ current waveform by calculating the time derivative of the fluorescence transient [17]. Specifically, we fitted the fluorescence transient with a 4-sigmoid fit and calculated its time-derivative as done in a previous study [18]. An example of this type of measurement is reported in Fig. 3a. As already reported by Hanemaaijer et al. [12], the kinetics of the Ca^2+^ signal associated with the AP varied with the distance from the soma with the onset of the Ca^2+^ transient in the distal axon preceding that in the proximal axon. This signal translated in a substantial delay between the peak of the Ca^2+^ current in the distal axon occurring before the peak of the somatic AP, and the peak of the Ca^2+^ in the proximal part of the AIS occurring after the peak of the somatic AP (Fig. 3b). This behaviour was consistently observed in N = 61 cells tested in this way (Fig. 3b), with a significant delay between the peaks of the distal and the proximal Ca^2+^ currents of 402 ± 177 μs (p<0.01, paired t-test). The evident anticipation of the AP peak, also with respect to the somatic AP peak, suggests that the Ca^2+^ influx associated with the AP is mainly mediated by VGNCs in the distal axon, whereas it is mainly mediated by VGCCs in the proximal axon. To address this hypothesis, we assessed the effect of locally delivering 80 nM hwtx on the Ca^2+^ influx and current associated with the AP in the AIS. In the example of Fig. 3c, hwtx substantially decreased the distal Ca^2+^ influx and current associated with the AP, whereas it only slightly decreased the proximal Ca^2+^ influx. To compare the effects of partially blocking Na_v_1.2 with those produced by blocking VGCCs, we first analysed the effects of inhibiting individual types of VGCCs. In detail, we blocked L-type VGCCs with 20 μM isradipine, P/Q-type VGCCs with 1 μM ω-agatoxin-IVA, N-type VGCCs with 1 μM ω-conotoxin-GVIA, R-type VGCCs with 1 μM of SNX482, and T-type VGCCs with 5 μM ML218 and 30 μM NNC550396. The results of this accurate analysis, reported in Supplementary Fig. S3, indicate that all types of VGCCs, at different extent, contribute to the axonal Ca^2+^ influx and current associated with the AP. Thus, to assess the effect of blocking all VGCCs, we locally delivered a cocktail with all inhibitors tested, at the same concentrations. In the example of Fig. 3d, the block of VGCCs substantially decreased the proximal Ca^2+^ influx and current associated with the AP, whereas the decrease of the distal Ca^2+^ current was only comparable to that produced by local delivery of 80 nM hwtx. We then compared the analysis of the change of the Ca^2+^ current peak in N = 8 cells where the effect of locally delivering hwtx was tested, and in N = 8 cells where the effect of locally delivering the VGCC blockers cocktail was tested (Fig. 3e). Whereas the partial block of Na_v_1.2 produced a marginal decrease of ∼10% of the Ca^2+^ current in the proximal AIS, compared to the significant decrease of >60% produced by blocking VGCCs (p<0.01, paired t-test), in the distal AIS hwtx and VGCCs blockers produced a similar and significant decrease of the Ca^2+^ current (p<0.01, paired t-test). In summary, the reduction of ∼26% of the distal Ca^2+^ current (Fig. 3e) and the block of 30-40% Na_v_1.2 produced by locally delivering 80 nM hwtx are consistent with a scenario where ∼70% of the Ca^2+^ influx associated with the AP, at distal sites of the AIS, is mediated by Na_v_1.2. The block of 30-40% Na_v_1.2 can therefore reduce the K^+^ current mediated by CAKCs widening the AP waveform at distal sites (Fig. 2).

**Fig. 3.**
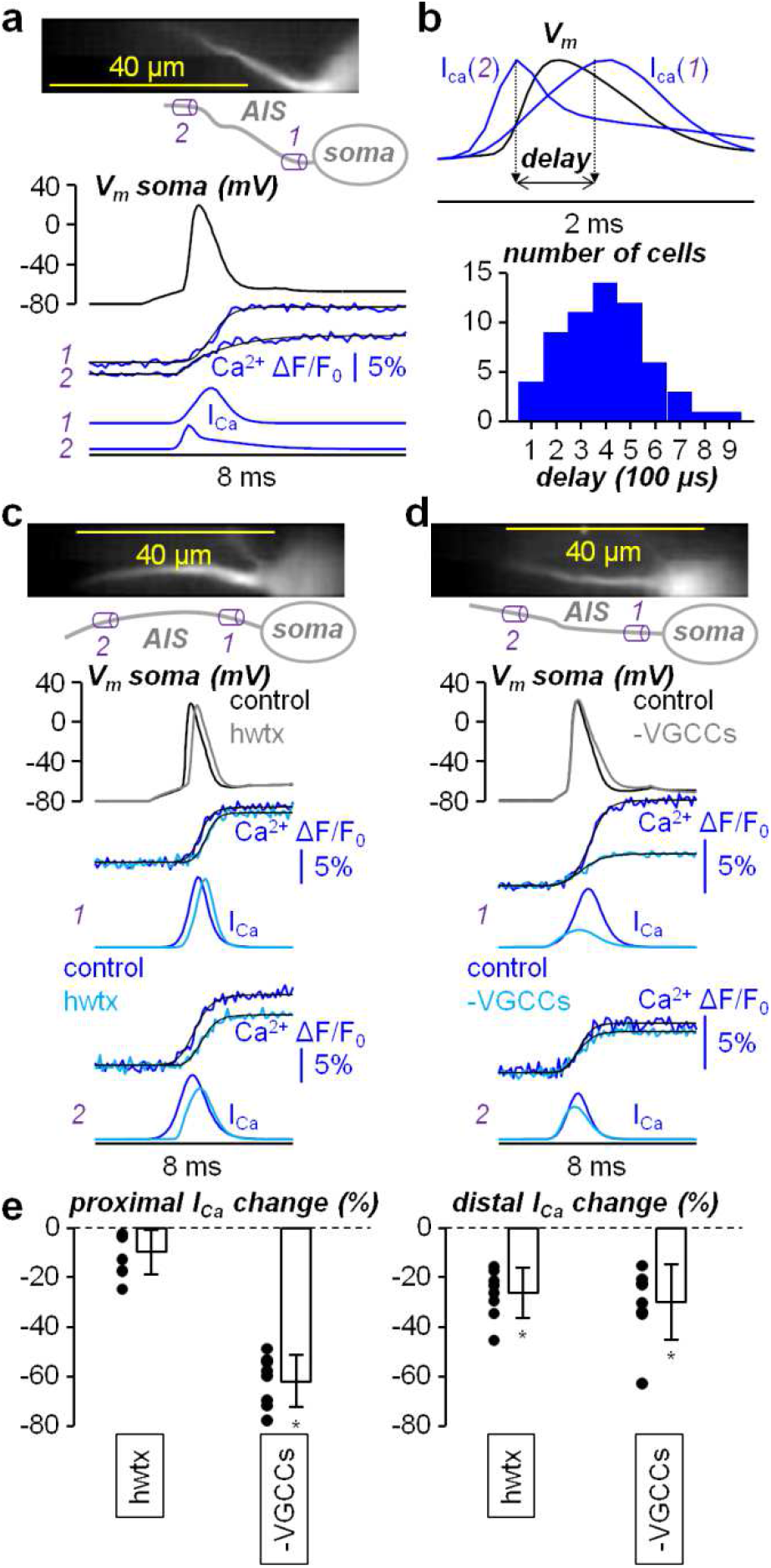
Effect of hwtx (block of Nav1.2) on the Ca^2+^ influx in the AIS. **a**, Top, fluorescence image (Ca^2+^ indicator OG5N) of the AIS and its reconstruction with a proximal region (*1*) and a distal region *(2)* indicated. Bottom, somatic AP and associated Ca^2+^ transients fitted with a 4-sigmoid (product of four sigmoids) function in *1* and *2*; the Ca^2+^ currents calculated as the time-derivative of the fits are reported below. **b**, Top, from the experiment in a, Ca^2+^ currents superimposed to the somatic AP. Bottom, over N = 61 cells tested, distribution of the delay between the distal and the proximal Ca^2+^ current peaks. **c**, Top, Ca^2+^ fluorescence image of the AIS and its reconstruction with a proximal region (*1*) and a distal (*2*) region indicated. Bottom, somatic AP and associated Ca^2+^ transients and currents in control solution and after local delivery of 80 nM of hwtx (top) *1* and *2*. **d**, In another cell, same protocol as that reported in panel c, but with local delivery of a cocktail blocking all VGCCs (-VGCCs) comprising 20 μM isradipine (L-type), 1 μM ω-agatoxin-IVA (P/Q-type), 1 μM ω-conotoxin-GVIA (N-type), 1 μM SNX482 (R-type), 5 μM ML218 + 30 μM NNC550396 (T-type). **e**, Left, single values and percentage change of the proximal Ca^2+^ current peak after addition of hwtx (N = 8 cells, mean ± SD = -9.7 ± 8.9) or after addition of VGCCs blockers (N = 8 cells, mean ± SD = -62.2 ± 10.5). Right, in the same cells, single values and percentage change of the distal Ca^2+^ current peak after addition of hwtx (mean ± SD = -26.2 ± 10.1) or after addition of VGCCs blockers (mean ± SD = -30.0 ± 15.1). “*” indicates a significant change (p<0.01, paired t-test).

### The block of BK CAKCs widens the AP in the AIS preventing the widening caused by blocking Na_v_1.2

In keeping with the results reported in Fig. 3, we next assessed the regulation of the AP waveform by individual CAKCs. In the example of Fig. 4a, locally delivering 1 μM of the SK CAKC inhibitor apamin widened the AP waveform in the soma and in the proximal and distal sites of the AIS. Visually, this widening was larger towards the late phase of the repolarisation. Similarly, in the example of Fig. 4b, locally delivering 1 μM of the BK CAKC inhibitor iberiotoxin also widened the somatic and axonal AP waveforms, but in this case the widening was observed earlier during the repolarisation. The widening behaviours produced by apamin or iberiotoxin were consistently observed in all cells tested. In contrast, the effects produced by 1 μM of the IK CAKC inhibitor tram-34 were variable. In the example of Supplementary Fig. S4a, locally delivering tram-34 did not change the AP waveforms in the soma and at all sites of the AIS, but in 4/8 cells tested in this way a widening behaviour like that produced by apamin was observed, as indicated by the analysis of the ∫V_m_ signal (see Fig. 2b) reported in Supplementary Fig. S4b. Since the effect of blocking IK channels was variable, we conducted a further analysis on SK and BK CAKCs only. To see whether a Ca^2+^ source activates a particular CAKC, one can block only this particular Ca^2+^ source and see whether this block prevents the effects produced by CAKCs. Thus, we first tested the combination of VGCCs as Ca^2+^ source by using the cocktail of blockers already used in the experiments of Fig. 3. In the example of Fig. 4c, we first blocked VGCCs, observing a widening of the AP waveform in the soma and in the proximal and distal sites of the AIS, and then added apamin to the VGCCs blockers cocktail. The block of SK CAKCs did not further change the somatic and axonal AP waveforms once VGCCs were blocked. In contrast, in the example of Fig. 4d, after observing a widening of the somatic and axonal AP waveforms produced by blocking VGCCs, addition of iberiotoxin caused a further widening of the V_m_ signals. The quantitative assessment of the effects of apamin and iberiotoxin was done in four separate groups of 7 cells in which the two inhibitors were tested first in control conditions and then after blocking VGCCs. In control conditions, the delivery of these inhibitors produced a significant increase of the ∫V_m_ signal (p<0.01, paired t-test), that was larger at distal sites of the AIS for iberiotoxin (Fig. 4d). The widening effects produced by apamin were similar to those produced by blocking VGCCs (N = 14 cells). When VGCCs were previously blocked, however, further addition of apamin did not change the ∫V_m_ signal (Fig. 4e). In contrast, further addition of iberiotoxin increased the ∫V_m_ signal, in a significant manner (p<0.01, paired t-test), by ∼12% at the proximal AIS sites and by ∼21% at the distal AIS sites (Fig. 4e). We concluded that SK CAKCs are activated exclusively by VGCCs, whereas BK CAKCs are also activated by other Ca^2+^ sources. Thus, to finally establish whether BK CAKCs are the target of Ca^2+^ entering through Na_v_1.2, we measured the change in the AP waveform produced by locally delivering 80 nM hwtx, in the presence of 1 μM iberiotoxin. In contrast to the cell of Fig. 2a-b where hwtx was applied in control solution, in the cell of Fig. 5a hwtx local delivery did not change the ∫V_m_ signals. In this final quantitative assessment performed in a group of 7 cells (Fig. 5b), addition of hwtx in the presence of iberiotoxin did not change the ∫V_m_ signal, in contrast to the increase of the ∫V_m_ signal observed without iberiotoxin. This result indicates that the block of BK CAKCs prevents the widening of the AP waveform, in the AIS, produced by the block of Na_v_1.2. We concluded that the Na_v_1.2 Ca^2+^ influx targets BK CAKCs, in this way shaping the AP waveform at its generating site.

**Fig. 4.**
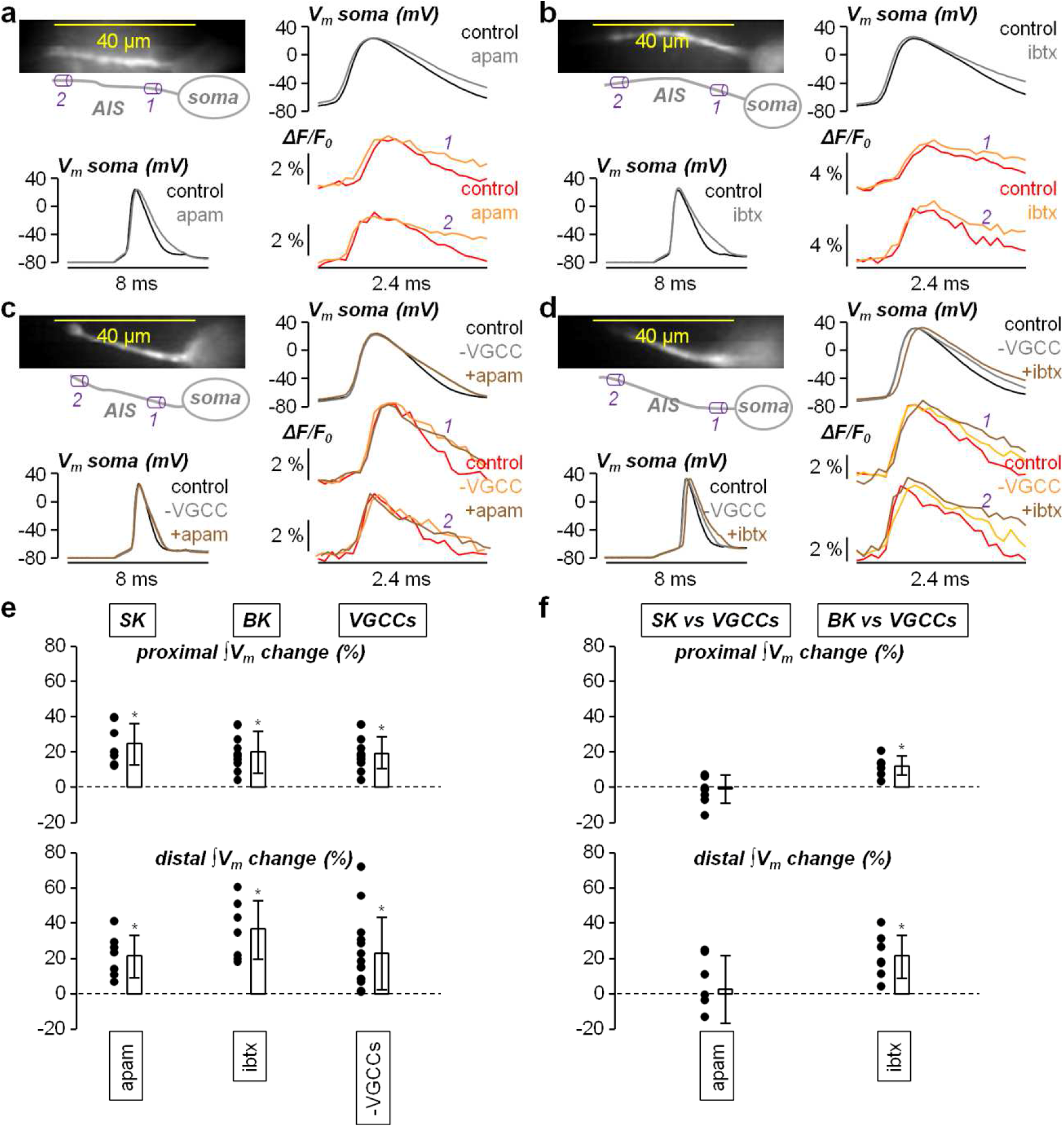
Effect of blocking SK CAKCs or BK CAKCs on the AP waveform in the AIS. **a**, Left, fluorescence image (JPW1114) of the AIS and its reconstruction with a proximal region (*1*) and a distal (*2*) region indicated; somatic AP before and after locally delivering 1 μM of the SK CAKC inhibitor apamin (apam) reported below. Right, somatic AP in control solution and after apamin delivery and associated V_m_ transients in *1* and *2*. **b**, In another cell, same protocol of panel a but by locally delivering 1 μM of the BK CAKC inhibitor iberiotoxin (ibtx). **c**, In another cell, same protocol of panel a but by sequentially blocking first VGCCs only and then VGCCs and SK CAKCs with additional apamin. **d**, In another cell, same protocol of panel a but by sequentially blocking first VGCCs only and then VGCCs and BK CAKCs with additional iberiotoxin. **e**, Top, single values and percentage change of the proximal ∫V_m_ signal maximum (see analysis in Fig. 2) after delivering apamin (N = 7 cells, mean ± SD = 24.7 ± 11.7), iberiotoxin (N = 7 cells, mean ± SD = 20.1 ± 11.9) or the cocktail of VGCCs blockers (N = 14 cells, mean ± SD = 19.0 ± 8.9). Bottom, in the same cells, single values and percentage change of the distal ∫V_m_ signal maximum after delivering apamin (mean ± SD = 21.7 ± 12.0), iberiotoxin (mean ± SD = 36.9 ± 16.6) or the cocktail of VGCCs blockers (mean ± SD = 22.9 ± 20.5). **f**, Top, with respect to the block of VGCCs, single values and percentage change of the proximal ∫V_m_ signal maximum after delivering apamin (N = 7 cells, mean ± SD = -2.1 ± 7.9) or iberiotoxin (N = 7 cells, mean ± SD = 12.1 ± 5.4). Bottom, in the same cells, single values and percentage change of the distal ∫V_m_ signal maximum after delivering apamin (mean ± SD = 2.4 ± 19.1) or iberiotoxin (mean ± SD = 21.5 ± 12.2). “*” indicates a significant change (p<0.01, paired t-test).

**Fig. 5.**
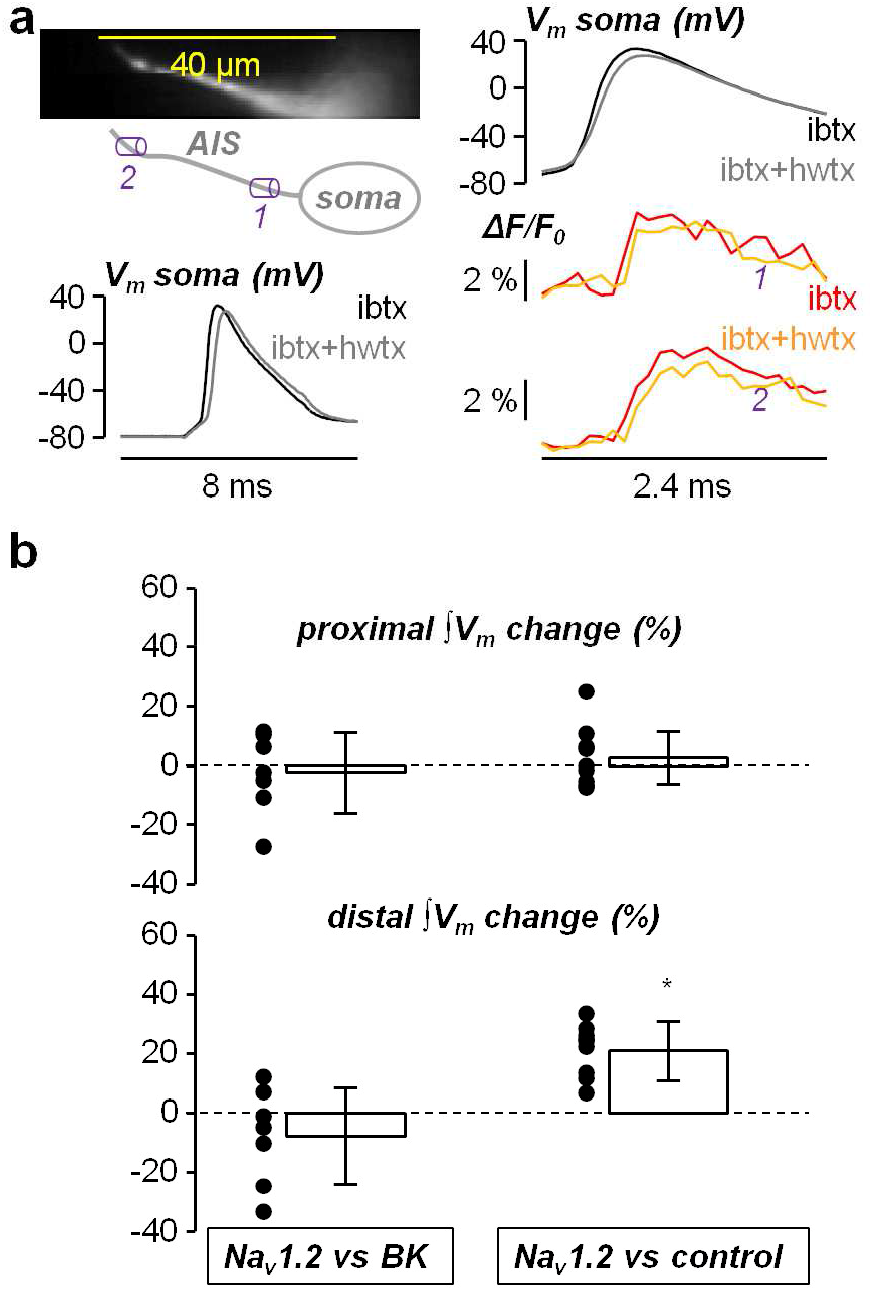
Effect of hwtx on the AP waveform in the AIS after blocking BK channels. **a**, Left, fluorescence image (JPW1114) of the AIS and its reconstruction with a proximal region (*1*) and a distal region (*2*) indicated; somatic AP before and after locally delivering 80 nM hwtx in the presence of iberiotoxin (1 μM) reported below. Right, somatic AP and associated V_m_ transients in *1* and *2* in the presence of iberiotoxin before and after hwtx delivery. **b**, Single cell values and percentage change (N=7 cells) of the ∫V_m_ signal maximum after locally delivering hwtx in the presence of iberiotoxin in proximal regions (mean ± SD = -2.3 ± 13.8), and in distal regions (mean ± SD = -7.9 ± 16.3). The statistics of the ∫V_m_ signal maximum after locally delivering hwtx in control solution (Fig. 2c) are also reported for comparison on the right. “*” indicates a significant change (p<0.01, paired t-test).

### Na_v_1.6 does not mediate Ca^2+^ influx associated with the AP in the AIS

The experiments reported in Figs. 1-5 demonstrate that Na_v_1.2 in the AIS shapes the AP waveform during the rising phase, by providing a large fraction of the inward current, and in the falling phase by regulating the Ca^2+^-activated outward current. The other VGNC (Na_v_1.6) also contributes to the AP shaping by providing the other Na^+^ inward current, but whether it also regulates the Ca^2+^-activated outward current remained to be established. In a final set of experiments, we addressed this question by performing the same tests we did with hwtx (Figs. 1-3), but in this case using the Na_v_1.6 inhibitor 4,9-anhydrotetrodotoxin (attx). This molecule exhibits moderate selectivity for Na_v_1.6 with respect to Na_v_1.2 as measured in HEK293 cells using automated patch clamp recordings (Supplementary Fig. S5). Specifically, at 10 nM and 33 nM, attx blocks 20 ± 13% and 30 ± 13% Na_v_1.6, respectively, whereas it blocks 9 ± 11% and 16 ± 12% Na_v_1.2 respectively. In AP tests performed in brain slices, attx at 400 nM did not affect the AP, whereas the smallest concentration at which we consistently observed a delay in the AP offset was 800 nM (Supplementary Fig. S5). At 1600 nM, attx consistently produced a stronger AP inhibition (Supplementary Fig. S5). Similarly to hwtx, the effects of attx at all concentrations were stable for several minutes after delivery. Thus, imaging experiments in brain slices were performed by using attx at the concentration of 800 nM providing the minimal perturbation of the somatic AP. Surprisingly, in the cell reported in Supplementary Fig. S6a, local delivery of 800 nM produced an increase of the distal Na^+^ transient, associated with a large increase of the slow non-inactivating component of the Na^+^ current (Supplementary Fig. S6b). A large increase of the Na^+^ transient and of the I_slow_ at distal sites was observed in 4/8 tested with this protocol (Supplementary Fig. S6c), suggesting that attx slows down the kinetics of inactivation of VGNCs. Consistently with a variable modulation of the I_slow_, attx also produced variable effects on the axonal AP waveform. While in the cell reported in Supplementary Fig. S7a-b the local delivery of attx did not change the AP waveform in the AIS, in the cell reported in Supplementary Fig. S7c-d attx widened the AP at the distal site of the AIS. In N = 7 cells tested with this protocol, the widening of the distal axonal AP was observed in 4 cells, whereas the AP waveforms did not change in 3 cells (Supplementary Fig. S7e). The complex interaction of attx with VGNCs does not allow the use of this tool to analyse in detail the role of Na_v_1.6. Nevertheless, since attx prolongs the Na^+^ influx (Supplementary Fig. S6a), the Ca^2+^ influx should be also prolonged if the channel affected by attx is permeable to Ca^2+^. In the cell of Supplementary Fig. S8a, however, the local delivery of attx did not change the axonal Ca^2+^ current and no significant change of the proximal or distal axonal Ca^2+^ current was measured in N = 7 cells tested with this protocol (Supplementary Fig. S8b). Since Na_v_1.2 is permeable to Ca^2+^ (Fig. 3), this result indicates that attx affects the kinetics of inactivation of Na_v_1.6 in murine brain slices and that this VGNC isoform is not contributing to the early Ca^2+^ influx. We concluded that only Na_v_1.2 mediates Ca^2+^ influx during the generating AP in the AIS and that only this isoform is functionally coupled to BK CAKCs.

### The Na_v_1.2-BK channels interaction is mimicked by simulations in a NEURON model

To mimic the experimental AP generation in the AIS, we built a NEURON model on the base of our dataset. Starting from the model reported in Hallermann et al. [23] and already utilised in our previous article [15], we adapted the AIS morphology to the mouse L5 pyramidal neuron and replaced the Na_v_1.2 and Na_v_1.6 channel models with those proposed in Mainen et al. [28], distributed consistently with the results reported by Hu et al. [7] (see Fig 6a). From this starting model, we modified the channel functions, as described in the Methods, in order to match the experimental Na^+^ and V_m_ signals in the AIS. We then introduced Ca^2+^ permeability to Na_v_1.2 corresponding to 0.4% with respect to the permeability for Na^+^ (as suggested by Hanemaaijer et al. [12]) and we introduced a low voltage-activated VGCC (LVAC) and a high voltage-activated VGCC (HVAC). We introduced an endogenous Ca^2+^ buffer and, when matching Ca^2+^ imaging experiments only, a Ca^2+^ buffer corresponding to 2 mM OG5N. Finally, we introduced SK and BK CAKCs and we explicitly introduced a coupling between SK channels and LVGCCs as described in the Methods. The relative spatial distributions of VGNCs, VGCCs, BK and SK channels, reported in Fig 6a, were set in order to reproduce the experimental results. Fig. 6b shows the proximal and distal V_m_ waveforms, the Na^+^ transients and the Ca^2+^ transients obtained by running computer simulations with the NEURON model. To validate the ability of the model to match the experimental evidence, we simulated the block of 80% VGCCs, of 80% SK channels (experiments with apamin) or of 80% BK channels (experiments with iberiotoxin). As shown in Fig 6c, computer simulations qualitatively replicate the experimental changes of the AP shape reported in Fig 4, indicating that the model realistically reproduces the activation of CAKCs during the AP generation. Thus, we eventually mimicked the effect of hwtx to produce a partial block of Na_v_1.2, that replicates the experimental shift of the AP with minimal change of the peak and the reduction of the global Na^+^ transient of ∼30% (see Fig.1). As shown in Fig 6d, the simulation reproduces the experimental reduction of the Na^+^ and Ca^2+^ transients and, more importantly, the widening of the distal AP (see Fig.2). Thus, the NEURON model that we built predicts the interplay between Na_v_1.2 and BK CAKCs associated with the generation of the AP in the AIS and replicates, in a computer simulation, the experimental widening of the axonal AP produced by hwtx delivery.

**Fig. 6.**
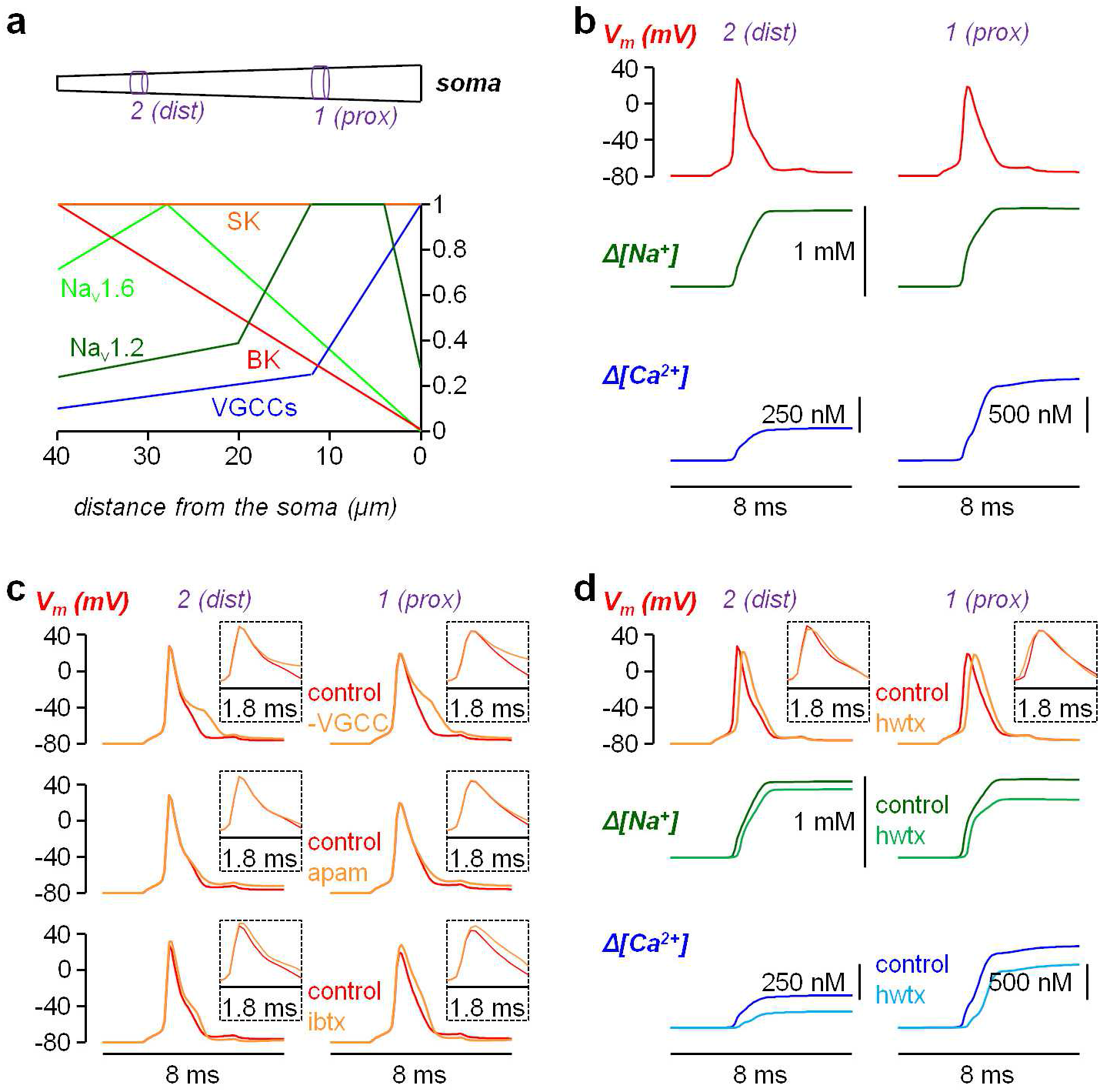
NEURON simulations of the AP generation in the AIS. **a**, Top, structure of the AIS (40 μm long) with proximal (*1*) compartment and distal (*2*) compartment at 10-12 μm and 30-32 μm from the soma, respectively. Bottom, normalised distributions of Na_v_1.2 VGNCs (dark green), Na_v_1.6 VGNCs (light green), VGCCs (blue), SK CAKCs (orange) and BK CAKCs (red) along the AIS. **b**, V_m_ (AP waveform), free Na^+^ and free Ca^2+^ from a computer simulation of the NEURON model in the proximal and distal regions. The presence of OG5N was taken into account for the free Ca^2+^ estimate. **c**, V_m_ (AP waveform) from computer simulations of the NEURON model in the proximal and distal regions in control conditions and after mimicking the delivery of VGCCs blockers (80% VGCCs reduction, top trace), of apamin (80% SK CAKCs reduction, middle trace) or of iberiotoxin (80% BK CAKCs reduction, bottom trace). The changes of the AP waveform are reported in the insets. The simulations qualitatively reproduce the experimental results reported in Fig. 4. **d**, V_m_ (AP waveform), free Na^+^ and free Ca^2+^ from computer simulations of the NEURON model in the proximal and distal regions in control conditions and after mimicking the delivery of hwtx. The presence of OG5N was taken into account for the free Ca^2+^ estimate. The changes of the AP waveform are reported in the insets. The simulations qualitatively reproduce the experimental results reported in Figs. 1-3.

## Discussion

If co-expression of multiple isoforms of VGNCs occurs in the mammalian AIS, does each isoform carry one or more specific roles in the generation and propagation of the AP? In pyramidal neurons, the different temporal and spatial patterns of expression of Na_v_1.2 and Na_v_1.6, but also the diverse biophysical properties underlying activation and inactivation, suggest that these two channels contribute differently to the generation/propagation of APs [7], as well as in the ability to mediate high or low frequency firing [8]. In this study we demonstrate that the exclusive permeability of Na_v_1.2 to Ca^2+^ [12] allows activation of BK CAKCs in the AIS. In this way, the interplay between Na_v_1.2, mediating the AP rise, and BK channels regulating the peak and the early phase of the AP repolarisation, shapes the AP waveform at the site of generation. To reach this important conclusion, we combined ultrafast imaging techniques with pharmacological analyses using selective channel blockers. The most important molecule used in this study was G^1^G^4^-huwentoxin-IV, mutated from huwentoxin-IV, a peptide from the Chinese bird spider *Haplopelma schmidti* [27]. Using this potent Na_v_1.2 inhibitor, sufficiently selective against Na_v_1.6, we could block, in a reliable manner, 30-40% of Na_v_1.2 in the AIS of L5 pyramidal neurons. This permitted the detailed investigation of Na^+^ currents, V_m_ waveforms and Ca^2+^ currents, associated with the generating AP, in the AIS. We then analysed the change in AP waveform produced by selectively blocking different types of CAKCs, eventually establishing the interaction between Na_v_1.2 and BK channels. Notably, several other channels contribute to the early process of generation and propagation of the AP. Thus, we created a NEURON model that adds to previous models developed for the AIS of L5 pyramidal neurons [7, 12, 20-24] and that reproduces our experimental results. Using this model, we could simulate the functional interaction between Na_v_1.2 and BK channels and mimic the experimental results obtained by blocking Na_v_1.2 with G^1^G^4^-huwentoxin-IV.

The process of AP generation starts with a sub-threshold depolarisation that spreads from the soma (or physiologically from the dendrites through the soma) to the AIS. The observation of an early subthreshold component of the Na^+^ influx [15] indicates that VGNCs also contribute to amplify the axonal depolarisation before the AP onset [29]. Whereas the threshold for the AP initiation is largely determined by Na_v_1.6 [30], our results point for a major contribution of Na_v_1.2 in the Na^+^ current during the rising phase of the AP, in particular at its peak occurring at the AP upstroke, even in the distal sites of the AIS where Na_v_1.6 expression is dominant. Importantly, an early Ca^2+^ current occurring with kinetics similar to that of the Na^+^ current allows Ca^2+^ binding to its targets before the AP reaches its peak. This is crucial since BK CAKCs require concomitant Ca^2+^ binding and depolarisation in order to open [31]. At the peak of the AP, BK channels contribute to the total K^+^ current that counterbalances the non-inactivated Na^+^ current, together with the diverse voltage-gated K^+^ channels expressed in the AIS [32]. Thus, a decrease of the Na_v_1.2 current is counterbalanced by a decrease of the BK current, resulting in widening of the early phase of AP repolarisation with a modest change of the AP peak. The interplay between Na_v_1.2 and BK channels is therefore a major determinant of the AP peak and a key regulator of the AP shape. After the peak, other channels contribute to the shape of the AP repolarisation, in particular VGCCs and SK CAKCs. Our results also provide evidence of the precise role of these channels in the shaping of the AP.

The shape of the AP at its site of origin is the starting point of the consequent physiological processes occurring at the other sites of the cells. First, the AP waveform, modulated during propagation along the axons, regulates neurotransmitter release and therefore synaptic transmission at synaptic terminals, in this way providing an analogue component of the digital information process [33]. Second, the AP waveforms regulate neuronal firing. Interestingly, in CA1 hippocampal pyramidal neurons, it has been shown that BK CAKCs facilitate high-frequency firing and cause early spike frequency adaptation [34]. The regulation of the firing frequency is crucial for the coupling among the different sites of the cell that are believed to underlie conscious processing [35]. Specifically, in L5 pyramidal neurons, high-frequency bursts of APs back-propagate along the apical dendrite reaching the tuft and provoking Ca^2+^ electrogenesis [36]. Finally, neurons with sharper AP kinetics have been correlated with human individuals exhibiting high IQ scores [37].

The result reported in this study is therefore a potential interpretation key in the investigation of the variety of neurological disorders associated with critical mutations of Na_v_1.2 [38,39] or BK channels [40]. Among the hundreds of mutations of the *SCN2A* gene, encoding Na_v_1.2, those classified as “gain of function” are typically associated with mild epilepsy [41], or with severe epileptic encephalopathies [42]. These phenotypes are generally explained by an enhancement of activity of glutamatergic neurons, despite Na_v_1.2 is also expressed in GABAergic interneurons [43]. The range of neurological disorders associated with other Na_v_1.2 mutations is far more complex, including autism and intellectual disability [38]. In parallel, heterogeneous combinations of disorders including seizure and intellectual disability are associated with mutations of KCNMA-1, the gene encoding BK channels [40]. Thus, the functional link between Na_v_1.2 and BK channels can represent an element to unravel the complexity of these channelopathies using murine models. In a heterozygote *SCN2A* knock-out mouse, haploinsufficiency of Na_v_1.2 is associated with an autistic-like phenotype attenuated with age [44], but in another SCN2A^+/-^ model the phenotype is also characterized by epileptic seizures [45] consistent with hyperexcitability of L5 pyramidal neurons [46]. More recently, a mouse model carrying the K1422A mutation in the *SCN2A* gene has been generated [47]. This specific mutation is characterised by an increase in Ca^2+^ permeability of Na_v_1.2 and a mix of phenotypes including rare seizures. This is an example where the result reported in the present study can represent an important interpretation key to link the cellular mechanisms to the behavioural dysfunction in the mouse model.

## Methods

### Selectivity tests of G^1^G^4^-huwentoxin-IV (hwtx) and of 4,9-Anhydrotetrodotoxin (attx)

Preliminary data on G^1^G^4^-huwentoxin-IV (hwtx), without the analysis on its selectivity, were previously reported [48].Tests of the selectivity of hwtx and attx were performed on HEK293 cells stably expressing either Na_V_1.2 or Na_V_1.6 channels as described in a previous report [49]. Automated patch-clamp recordings in whole-cell configuration were performed using the SyncroPatch 384PE from Nanion (München, Germany). Chips with single-hole medium resistance of 4.52 ± 0.08 MΩ (N = 384 experiments) were used for recordings. Pulse generation and data collection were performed with the PatchControl384 v1.5.2 software (Nanion) and the Biomek v1.0 interface (Beckman Coulter). The intracellular solution contained (in mM): 10 CsCl, 110 CsF, 10 NaCl, 10 EGTA and 10 HEPES (pH 7.2, osmolarity 280 mOsm). The extracellular solution contained (in mM): 140 NaCl, 4 KCl, 2 CaCl_2_, 1 MgCl_2_, 5 glucose and 10 HEPES (pH 7.4, osmolarity 298 mOsm). Voltage clamp experiments were performed at a holding potential of −100 mV at room temperature (18-22°C) and currents were sampled at 20 kHz. Each peptide was prepared in the extracellular solution supplemented with 3% BSA and a single peptide concentration was applied to each cell. For establishing dose-response curves, the compounds were tested at a test potential of 0 mV for 50 ms with a pulse every 5 sec. The percentage of current inhibition by the peptides was measured at equilibrium of blockage or at the end of a 14-min application time. Notice that the peptide used in this study (hwtx) was different from the active toxin in the caged compound described in our previous publication [49].

### Brain slices solution, preparation and maintenance

Experiments in brain slices were performed at the Laboratory of Interdisciplinary Physics in Grenoble in accordance with European Directives 2010/63/UE on the care, welfare and treatment of animals. Procedures were reviewed by the ethics committee affiliated to the animal facility of the university (D3842110001). The extracellular solution that we used contained (in mM): 125 NaCl, 26 NaHCO_3_, 1 MgSO_4_, 3 KCl, 1 NaH_2_PO_4_, 2 CaCl_2_ and 20 glucose, bubbled with 95% O_2_ and 5% CO_2_. Mice (C57BL/6j, 21-35 postnatal days old) purchased from Janvier Labs (Le Genest-Saint-Isle, France) were anesthetised by isofluorane inhalation and the entire brain was removed after decapitation. Brain slices (350 μm thick) were prepared using the procedure described in a previous report [15], using a Leica VT1200 vibratome (Wetzlar, Germany). Slices were incubated at 37°C for 45 minutes and maintained at room temperature before use.

### Electrophysiology and imaging in brain slices

Slices with L5 pyramidal neurons in the somato-sensory cortex having an axon parallel to the slice surface were transferred to the recording chamber. Patch-clamp recordings were made at the temperature of 32-34°C using a Multiclamp 700A (Molecular Devices, Sannyvale, CA). The intracellular solution contained (in mM): 125 KMeSO_4_, 5 KCl, 8 MgSO_4_, 5 Na_2_-ATP, 0.3 Tris-GTP, 12 Tris-Phosphocreatine, 20 HEPES, adjusted to pH 7.35 with KOH. In Na^+^ imaging experiments, the Na^+^ indicator ING-2 (IonBiosciences, San Marcos, TX) was added to the intracellular solution at the concentration of 0.5 mM and recordings were started 20 minutes after achieving the whole cell configuration. In V_m_ imaging experiments, cell membranes were loaded with the voltage-sensitive dye D-2-ANEPEQ (JPW1114, 0.2 mg/mL, Thermo Fisher Scientific) for 30 minutes using a first patch clamp recording and then re-patched a second time with dye free solution. Finally, in Ca^2+^ imaging experiments, the Ca^2+^ indicator Oregon Green BAPTA-5N (Thermo Fisher Scientific) was added to the intracellular solution at the concentration of 2 mM and recordings were started 20 minutes after achieving the whole cell configuration. Somatic APs were elicited in current clamp mode by short current pulses of 1.5-2.5 nA through the patch pipette electrical and electrical signals were acquired at 20 kHz. The bridge was corrected offline by using the recorded injected current and the measured V_m_ was corrected for -11 mV junction potential. Na^+^ and Ca^2+^ imaging experiments and most of V_m_ imaging experiments were performed using an imaging system described in a previous report [15], based on an upright Scientifica SliceScope microscope equipped with a motorised XY translation stage and PatchStar manipulators (Scientifica, Uckfield, UK), and a 60X Olympus water immersion objective (NA = 1). Na^+^ and V_m_ indicators were excited using the 520 nm line of a LaserBank (Cairn Research, Faversham, UK) band-pass filtered at 517 ± 10 nm and directed to the preparation using a 538 nm long-pass dichroic mirror. The Ca^2+^ indicator was excited using the 465 nm line of the LaserBank, band-pass filtered at 469 ± 17 nm and directed to the preparation using a 495 nm long-pass dichroic mirror. Na^+^ and V_m_ fluorescence emission signals were band-pass filtered at 559 ± 17 nm or long-pass filtered at >610 nm respectively. Ca^2+^ fluorescence emission was band-pass filtered at 525 ± 25 nm. The size of the illumination spot, obtained using a custom-made telescope, was ∼30 μm. Image sequences were demagnified by 0.5X and acquired with a DaVinci 2K CMOS camera (SciMeasure, Decatur, GA) at 10 kHz with a pixel resolution of 500 nm in regions of 30×128 pixels. The other V_m_ imaging experiments were performed using an imaging system described in a previous report [25]. Fluorescence was excited using the 530 nm line of LDI-7 laser (89 North, Williston, VT), band-pass filtered at 531 ± 40 nm and directed to the preparation using a 575 nm long-pass dichroic mirror. Fluorescence emission was long-pass filtered at >610 nm and acquired with a NeuroCCD camera (Redshirt Imaging, Decatur, GA) at 20 kHz with a pixel resolution of ∼2 μm in regions of 4×26 pixels. In all experiments, image sequences were acquired for 8 ms.

### Pharmacology

All molecules were initially dissolved in stock solutions and small aliquots were kept at -20 °C before use. G^1^G^4^-huwentoxin-IV (hwtx) was synthesised by Smartox Biotechnology (Saint Egrève, France). ω-agatoxin-IVA (agaIVA), ω-conotoxin-GVIA (conGVIA), snx482, apamin (apam) and iberiotoxin (ibtx) were also purchased from Smartox Biotechnology. 4,9-Anhydrotetrodotoxin (attx) was purchased from Tocris (Bristol, UK). Tram-34 (tram) was purchased from HelloBio (Bristol, UK). All these molecules were initially dissolved in water at 100 μM concentration and then diluted in external solution at the desired concentration. 4-(2,1,3-Benzoxadiazol-4-yl)-1,4-dihydro-2,6-dimethyl-3,5-pyridinecarboxylic acid methyl 1-methylethyl ester (isradipine or isr) (HelloBio) was initially dissolved at 20 mM in DMSO and then diluted in external solution 20 μM just before use. This final solution was kept for a maximum of 1h to prevent the loss of dissolved molecules due to precipitation. 3,5-dichloro-*N*-[[(1α,5α,6-exo,6α)-3-(3,3-dimethylbutyl)-3-azabicyclo[3.1.0]hex-6-yl]methyl]-benzamide-hydrochloride (ML218 or ml) was purchased from TOCRIS and initially dissolved at 50 mM in DMSO. (1*S*,2*S*)-2-[2-[[3-(1*H*-Benzimidazol-2-yl)propyl]methylamino]ethyl]-6-fluoro-1,2,3,4-tetrahydro-1-(1-methylethyl)-2-naphthalenyl-cyclopropanecarboxylate-dihydrochloride (NNC550396 or nnc) was purchased from TOCRIS and initially dissolved at 25 mM in water. These two molecules were diluted together in external solution at the desired concentrations. The protocol to locally deliver the channel blockers is illustrated in Supplementary Fig. S2a.

### Data analysis

All data, from averages of 4 trials with identical somatic response, were analysed using custom-written code written in MATLAB. Initially, recordings including an AP were corrected for photo-bleaching using multi-exponential fits of fluorescence transients in single trials without an AP. Then, the fractional change of fluorescence from the first image (ΔF/F_0_) was calculated over the mean fluorescence in 5 μm regions at proximal and distal locations of the AIS (Supplementary Fig. S2b). For Na^+^ imaging experiments, the Na^+^ current was calculated as described in detail in a previous report [15]. Specifically, ΔF/F_0_ signals were initially converted into Δ[Na^+^] signals and then into a Na^+^ (or a positive charge) superficial density using an estimate of the diameter of the axon. The signal was fitted with a model function (see [15]) and the current surface density (or simply the current) was finally obtained by calculating the time derivative of this fit. In the analysis, a fast (I_fast_) and a slow (I_slow_) of the Na^+^ current were extracted (see Fig. 1b). For V_m_ imaging experiments, to unambiguously quantify a change in AP beyond the noise of the optical recording, we calculated the integral of the ΔF/F_0_ signal over a time window of 2.4 ms comprising the AP. For Ca^2+^ imaging experiments, the Ca^2+^ current was calculated as described in detail in a previous report [18]. Specifically, ΔF/F_0_ signals were fitted with a model function consisting on the product of four sigmoid functions and the current was obtained by calculating the time derivative of this fit.

### Statistics

The effect of a channel blocker on a specific signal was quantified by computing its percentage change following local delivery of the blocker with respect to the control condition. To assess the consistency of the effect, for each protocol and in at least seven cells and we performed a paired t-test comparing the values in control conditions and the values after local delivery of the channels blocker. We considered 0.01 as the threshold to describe an effect as significant.

### NEURON model

The NEURON model, built from the starting model reported in Hallermann et al. [23] (adapted for the mouse) and already utilised in our previous article [15], was developed by replicating the results of the computer simulations with V_m_, Na^+^ and Ca^2+^ imaging results from three cells where the somatic AP had very similar shape. The AIS (40 μm long) has semi-conical shape with initial diameter of 4.22 μm and final diameter of 1.73 μm and it is divided in 20 compartments of 2 μm length. We took the 6^th^ compartment (10-12 μm from the soma) as representative of the proximal AIS and the 16^th^ compartment (30-32 μm from the soma) as representative of the distal AIS. The construction of the new model, from the initial basic model, was achieved through a strategy consisting in a sequence of steps.

1. First, we replaced the two generic VGNCs with the Na_v_1.2 and Na_v_1.6 channel models proposed in Mainen et al. [28], imposing the distribution of the initial model also consistent with the results reported in Hu et al. [7]. The functions and distributions of both channels were then optimised in order to take into account the different activation and inactivation kinetics to match the experimental Na^+^ transients both in the proximal and in the distal parts of the AIS.
2. Second, we introduced Ca^2+^ permeability to Na_v_1.2 corresponding to 0.4% with respect to the permeability for Na^+^ (as suggested by Hanemaaijer et al. [12]), a LVAC, corresponding to T-type channels, and a HVAC, corresponding to the ensemble of L-type, P/Q-type, N-type and R-type channels. The two models were adopted from a model in Almog and Korngreen [50]. The channel distribution and density were set to match the results of the Ca^2+^ imaging experiment. The distribution of VGCCs was also consistent with the results reported by another laboratory [51]. In the simulations used to match experimental Ca^2+^ transients, we introduced a generic mobile Ca^2+^ buffer with standard kinetics [52] to mimic 2 mM OG5N in the patch pipette. In all simulations, we accounted for an immobile endogenous buffer with capacity of ∼40 in the distal AIS, consistent with estimates at presynaptic sites [53]. Finally, for Ca^2+^ extrusion and Ca^2+^ radial/longitudinal diffusion, we used the model proposed in Kim et al. [54].
3. Third, we modified the K^+^ channels from the initial model in the following manner. We introduced the inward rectifier K^+^ channels from Migliore at al. [55], the BK CAKS from Ait Ouares et al [25], the SK CAKC from Mahapatra et al. [56], and we reduced the original K+ channels in order to preserve the kinetics of the AP in the AIS under control conditions. In addition, we explicitly made SK channels activation dependent on Ca^2+^ entering from LVGCCs. These modified profile of K^+^ channels produced AP waveforms qualitatively consistent with the experimental results shown in Fig.4, when 80% of VGCCs, BK channels or SK channels were eliminated from the model (see simulations in Fig.6c).

With this model, we finally mimicked the delivery of 80 nM hwtx by changing the Na_v_1.2 function in order to obtain a reduction of the Na^+^ transients in the proximal and distal AIS consistent with the experiments reported in Fig.1. This was done by a full block of a non inactivating component of the Na_v_1.2 current, a 20% reduction of a fast inactivating component of the Na_v_1.2 current, a 6% reduction of the activation slope of the Na_v_1.2 current and a 50% reduction of the Ca^2+^ permeability.

## Data availability

The complete dataset of imaging/electrophysiological experiments in brain slices, used in this study, is available in the public repository Zenodo (doi: 10.5281/zenodo.5835995). The Neuron model, including the changes of parameters replicating the experiments with hwtx are available in the NeuronDB database at: http://modeldb.yale.edu/267355.

## Code availability

Matlab codes used for data analysis are available at https://github.com/MarcoCanepari/NAV12-BK-paper.

## Acknowledgements

This work was supported by the *Agence Nationale de la Recherche* through three grants (ANR-18-CE19-0024 - OptChemCom; Labex *Ion Channels Science and Therapeutics:* program number ANR-11-LABX-0015 and National Infrastructure France Life Imaging “Noeud Grenoblois”) and by the *Federation pour la Recherché sur le Cerveau* (FRC – Grant *Espoir en tête*, Rotary France). We are indebted to Fondation Leducq and the FEDER resources for financing the automated patch clamp setup.

## Supplementary information

**Supplementary Figure S1.**
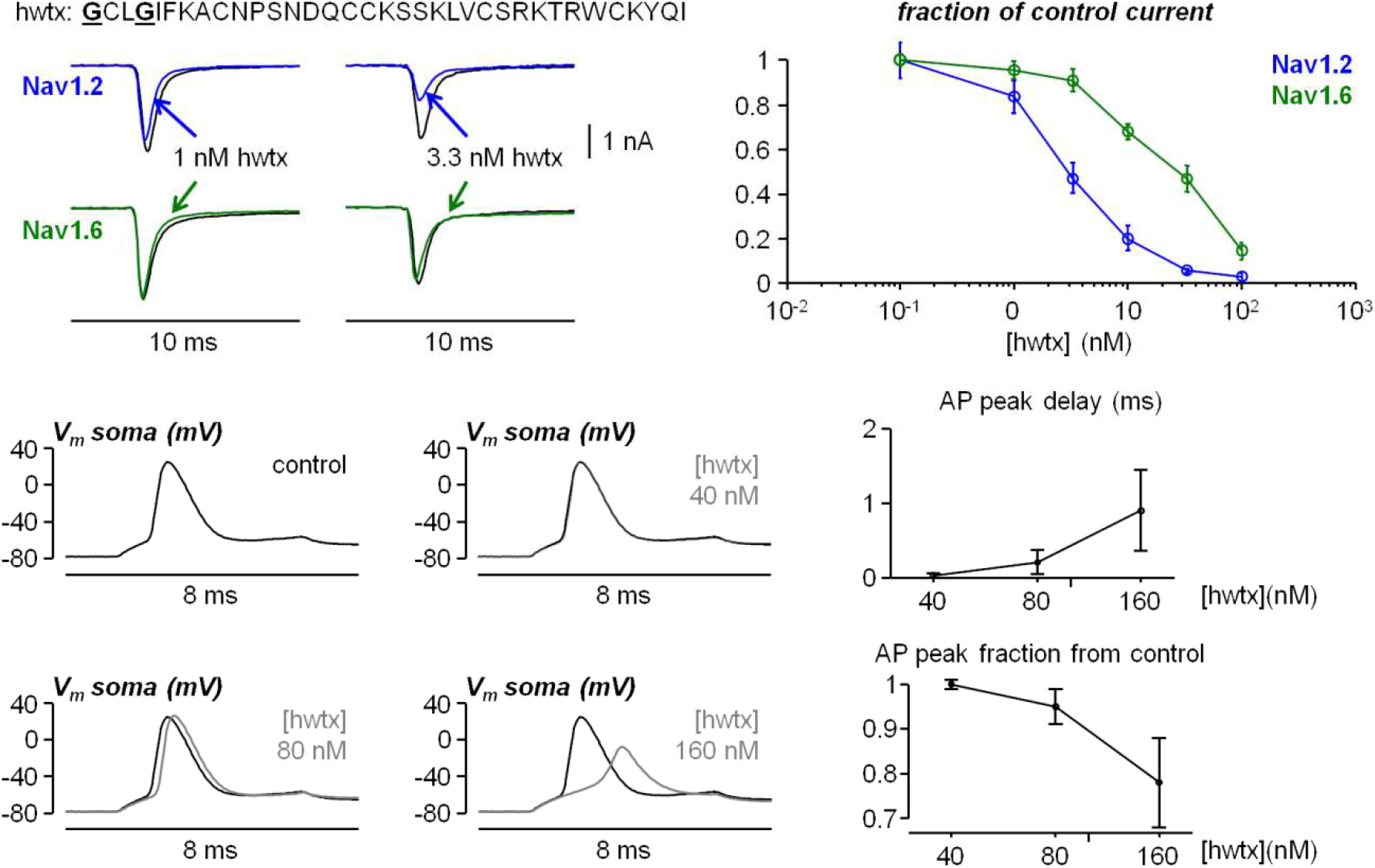
Selectivity of 1G4GHuwentoxin-IV (hwtx) for Nav1.2 against Nav1.6. Top-left, currents recorded in HEK293 cells expressing either Nav1.2 (blue) or Nav1.6 (green) in control solution and after addition of hwtx at 1 nM (left traces) or at 3.3 nM (right traces); the 35-aminoacids sequence of hwtx is reported; the two residues mutated from the wild type peptide are indicated with bold-underlined characters. Top-Right, fraction of Nav1.2 (green) or Nav1.6 (blue) current peak from control condition (no hwtx) against hwtx concentration (mean ± SD); each concentration-point has been calculated over 10-15 cells. Bottom-Left, in a L5 pyramidal neuron in brain slices, AP elicited in control condition (no hwtx) or in the presence of 40 nM, 80 nM or 160 nm hwtx. Bottom-Right, delay of the AP peak and AP peak fraction from control condition against hwtx concentration (mean ± SD, N = 7 cells).

**Supplementary Figure S2.**
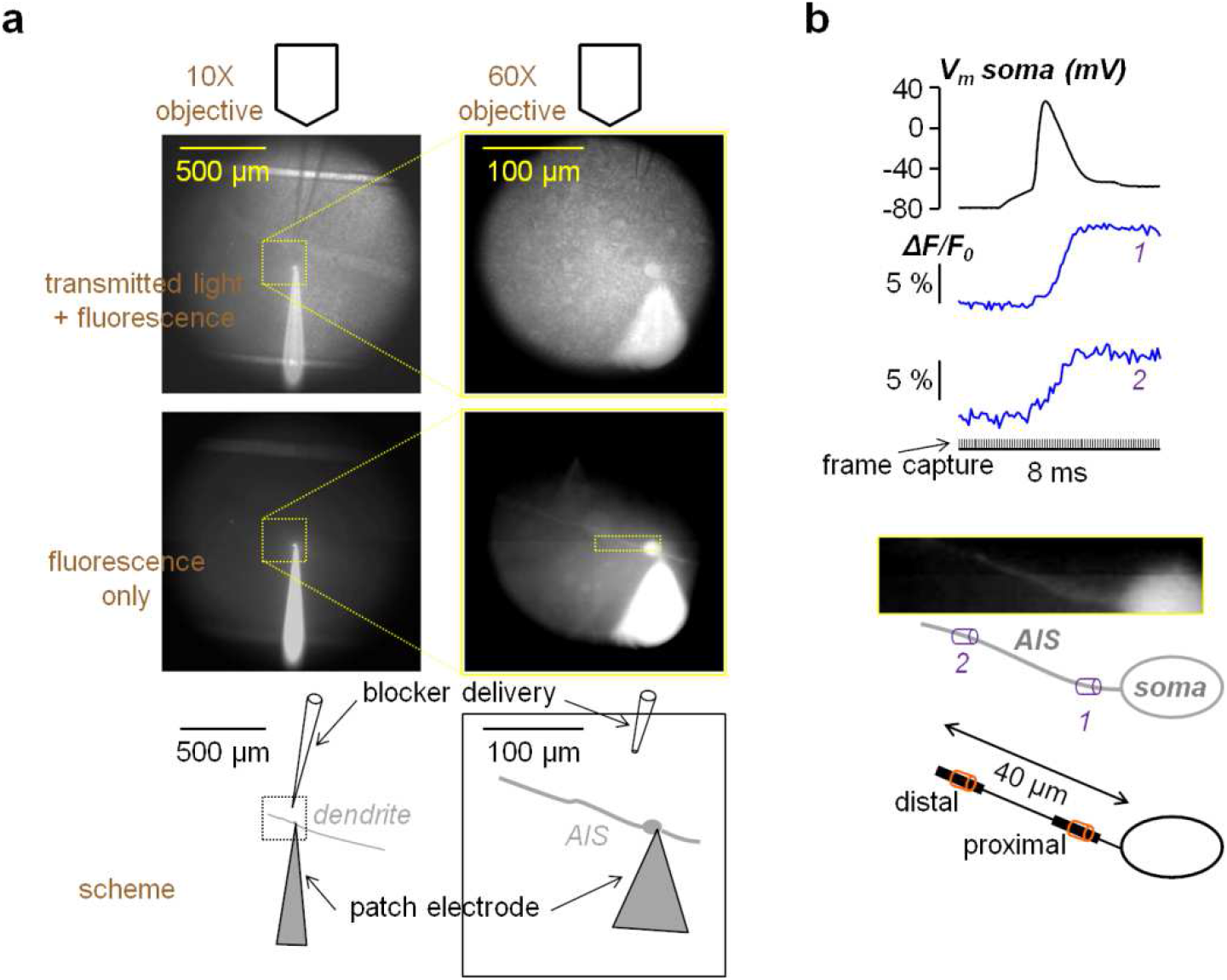
Illustrations of the experimental configuration and of the recording protocol used in the study. **a**, Illustration of the experimental configuration. A brain slice (somatosensory cortex) is visualised by the DaVinci-2K camera using a 10X magnification objective (images on the left) and a 60X magnification objective (images on the right); the images on the top are taken with transmitted IR light and fluorescence simultaneously; the images below are taken with fluorescence only; as indicated in the scheme below, a patch clamp electrode (also used to elicit APs and to record somatic V_m_) is used to load through the soma a L5 pyramidal neuron with a fluorescent indicator, while a pipette positioned near the AIS is used to locally deliver a solution containing a channel blocker, by gentle pressure application, for 1 minute. **b**, In the recording protocol, images of ∼64×15 μm^2^ comprising the AIS are acquired at the frame rate of 10 kHz; to unambiguously distinguish signals from proximal and distal areas of the AIS, with respect to the soma, we systematically analysed every signal in a region of 5 μm long at a position “*1*”, within 5 and 15 μm from the soma and in another region of 5 μm long at a position “*2*”, within 30 and 40 μm from the soma; on the top, an AP is elicited and recorded at the soma (at 20 kHz) and the fluorescence signals (Ca^2+^ in this example) at positions *1* and *2* are reported.

**Supplementary Figure S3.**
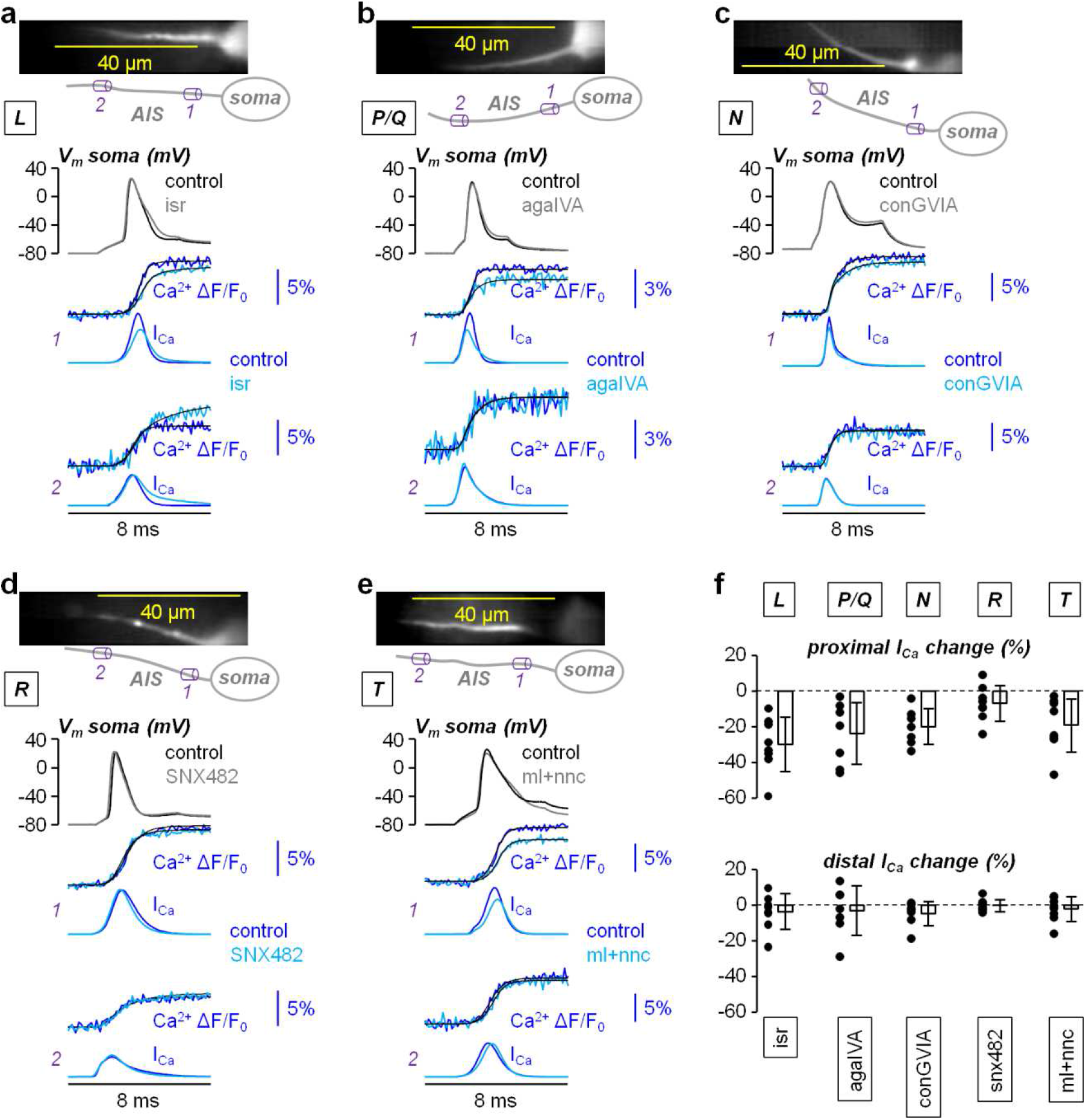
Analysis of the Ca^2+^ current (I_Ca_) components mediated by individual VGCC types in the AIS. **a**, Top, fluorescence image (Ca^2+^ indicator OG5N) of the AIS and its reconstruction with a proximal region (*1*) and a distal (*2*) region indicated. Middle, Somatic AP in control solution and after locally delivering 20 μM of the L-type VGCC inhibitor isradipine (isr). Bottom, Ca^2+^ transients associated with the APs above fitted with the product of four sigmoid functions (black traces) and I_Ca_ signals derived from the fits. **b**, Same protocol of panel a, but locally delivering 1 μM of the P/Q-type VGCC inhibitor ω-agatoxin-IVA (agaIVA). **c**, Same protocol of panel a, but locally delivering 1 μM of the N-type VGCC inhibitor ω-conotoxin-GVIA (conGVIA). **d**, Same protocol of panel a, but locally delivering 1 μM of the R-type VGCC inhibitor SNX482. **e**, Same protocol of panel a, but locally delivering 5 μM and 30 μM, respectively, of the T-type VGCC inhibitors ML218 and NNC550396 (ml+nnc). **f**, Top, single values and percentage change of the I_Ca_ peak in proximal regions after addition of isr (N = 7 cells, mean ± SD = -29.9 ± 17.0), after addition of agaIVA (N = 8 cells, mean ± SD = -23.9 ± 17.4), after addition of conGVIA (N = 7 cells, mean ± SD = -20.2±10.1), after addition of SNX482 (N = 8 cells, mean ± SD = -7.1 ± 10.0) or after addition of ml+nnc (N = 8 cells, mean ± SD = -19.0 ± 14.9). Bottom, in the same cells, single values and percentage change of the I_Ca_ peak in distal regions after addition of isr (mean ± SD = -3.8 ± 10.0), after addition of agaIVA (mean ± SD = -3.3 ± 13.9), after addition of conGVIA (mean ± SD = -4.8 ± 6.8), after addition of SNX482 (mean ± SD = -0.2 ± 3.4) or after addition of ml+nnc (mean ± SD = -2.3 ± 7.0).

**Supplementary Figure S4.**
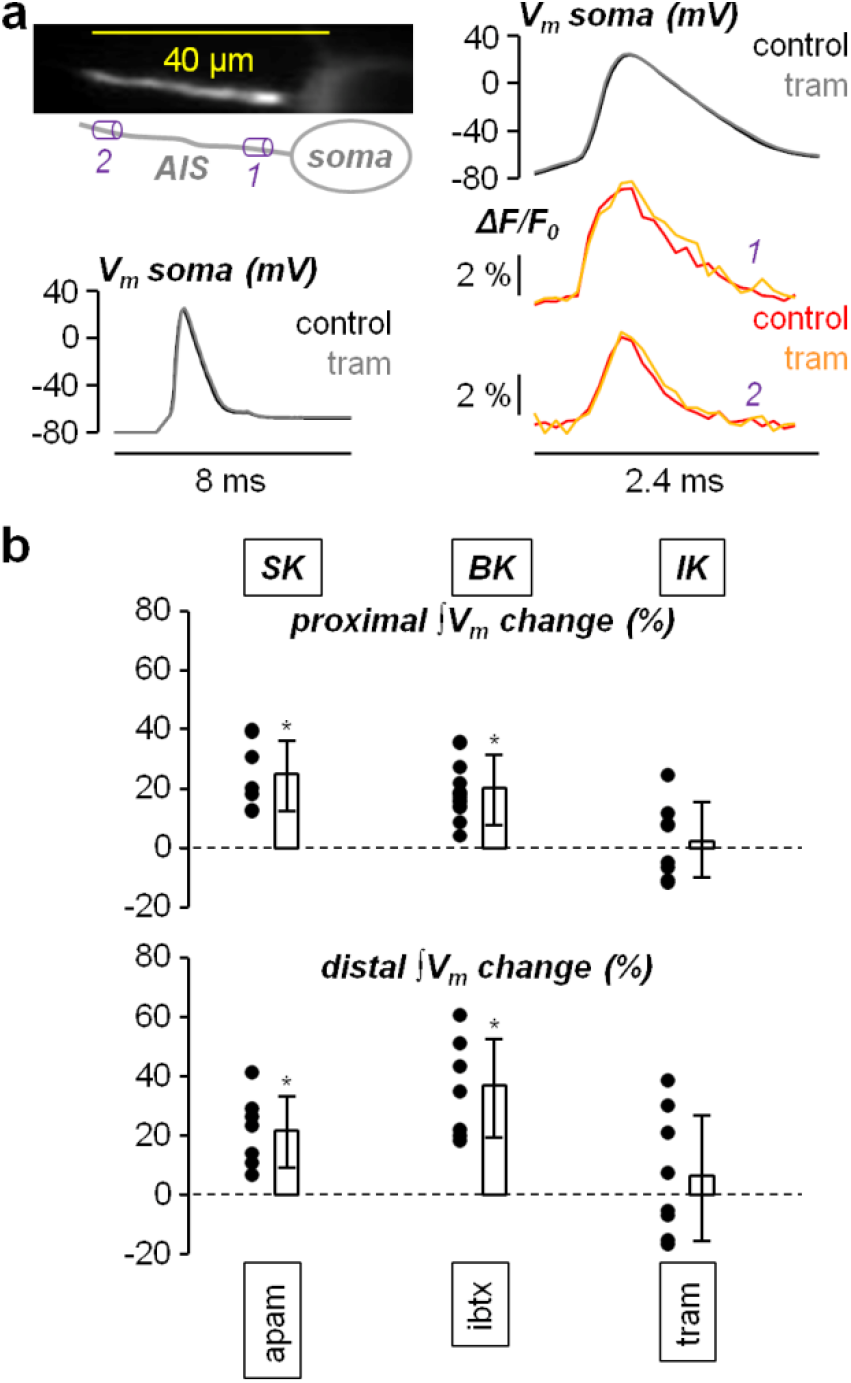
Analysis of the effect of blocking IK CAKCs on the AP waveform in the AIS. **a**, Left, fluorescence image (voltage sensitive dye JPW1114) of the AIS and its reconstruction with a proximal region (*1*) and a distal (*2*) region indicated; a somatic AP before and after locally delivering 1 μM of the IK CAKC inhibitor tram-34 (tram) is reported below. Right, Somatic AP in control solution and after tram-34 delivery and associated V_m_ transients in *1* and *2*. **b**, Top, single values and percentage change of the proximal ∫V_m_ signal maximum (see analysis in Fig. 2 and in Fig. 4) after delivering tram-34 (N = 8 cells, mean ± SD = 2.2 ± 12.7); the values and statistics for the same test performed with the SK CAKC inhibitor apamin (apam) or with the BK CAKC inhibitor iberiotoxin (ibtx) (same as in Fig. 4a) are reported on the left. Bottom, in the same cells, single values and percentage change of the distal ∫V_m_ signal maximum after delivering tram-34 (mean ± SD = 6.7 ± 21.1); the values and statistics for the same test performed with the SK CAKC inhibitor apamin or with the BK CAKC inhibitor iberiotoxin (same as in Fig. 4a) are reported on the left.

**Supplementary Figure S5.**
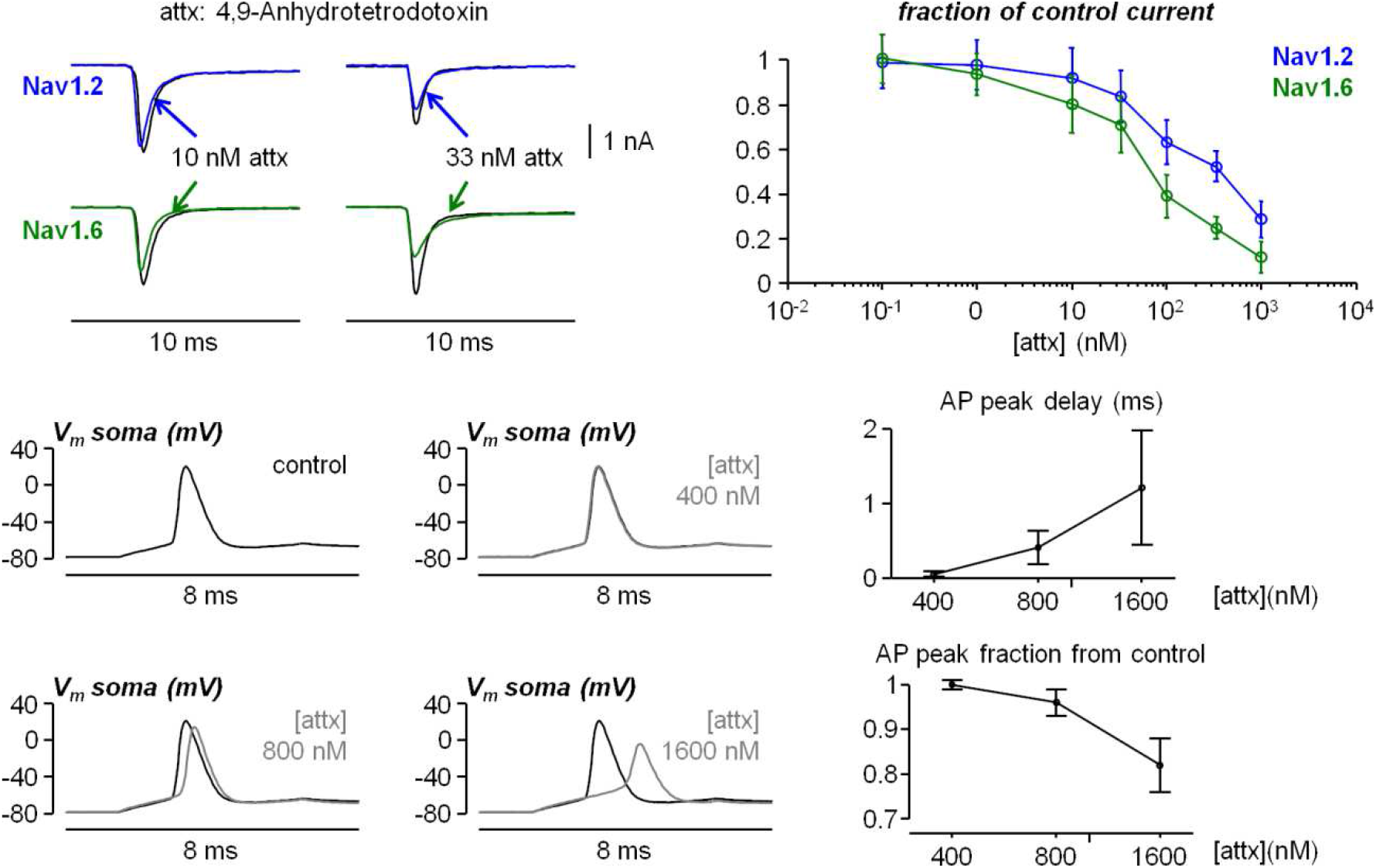
Selectivity of 4,9-Anhydrotetrodotoxin (attx) for Nav1.6 against Nav1.2. Top-left, currents recorded in HEK293 cells expressing either Nav1.2 (blue) or Nav1.6 (green) in control solution and after addition of attx at 10 nM (left traces) or at 33 nM (right traces). Top-Right, fraction of Nav1.2 (green) or Nav1.6 (blue) current peak from control condition (no attx) against attx concentration (mean ± SD); each concentration-point has been calculated over 10-15 cells. Bottom-Left, in a L5 pyramidal neuron in brain slices, AP elicited in control condition (no attx) or in the presence of 40 nM, 80 nM or 160 nm attx. Bottom-Right, delay of the AP peak and AP peak fraction from control condition against attx concentration (mean ± SD, N = 7 cells).

**Supplementary Figure S6.**
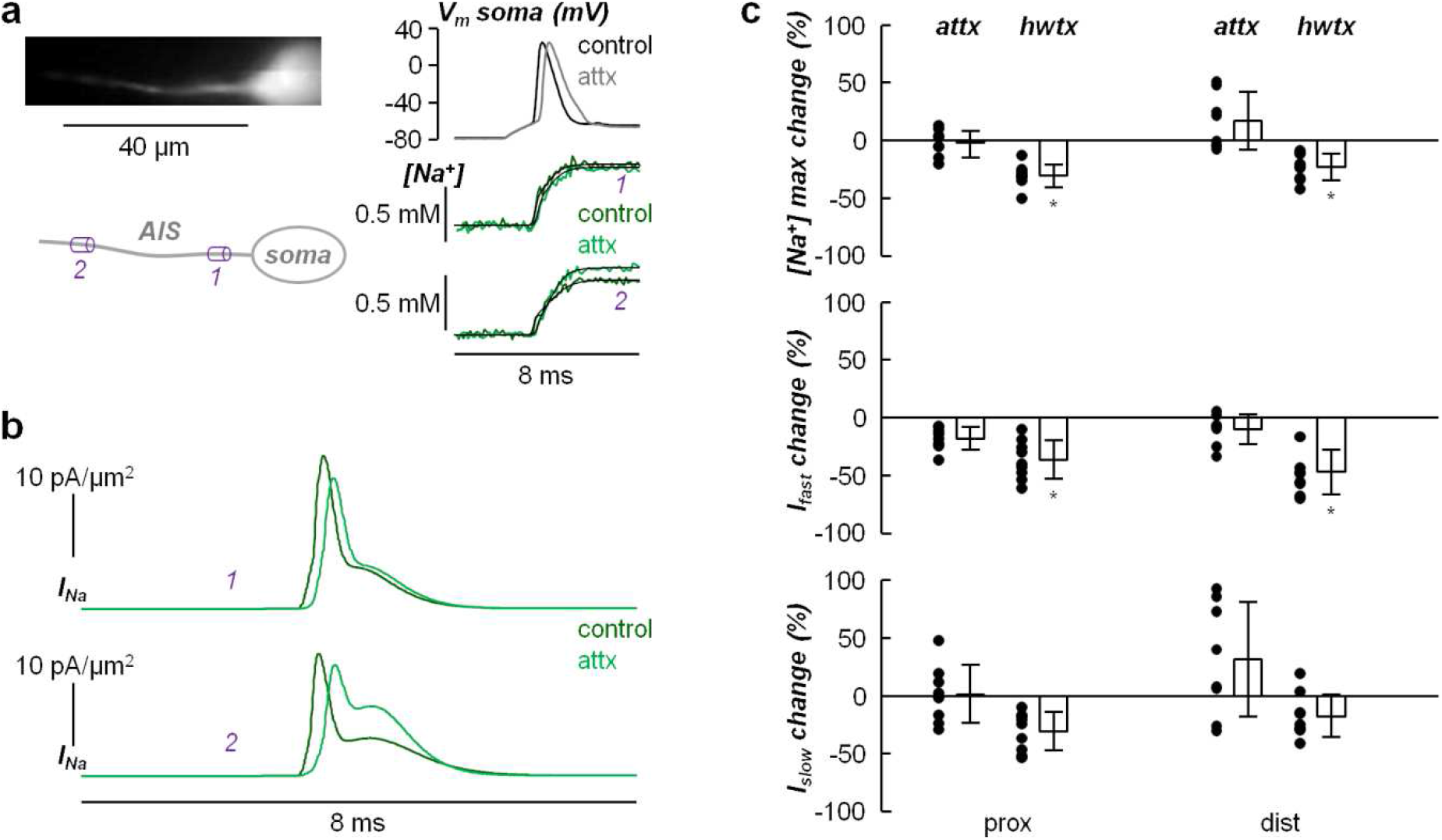
Analysis of the effect of 4,9-anhydrotetrodotoxin (attx) on the Na^+^ influx in the AIS. **a**, Left, fluorescence image (Na^+^ indicator ING-2) of the AIS and its reconstruction with a proximal region (*1*) and a distal (*2*) region indicated. Right, Somatic AP in control solution and after locally delivering 800 nM of attx (top) and associated corrected Na^+^ transients fitted with a model function in *1* and *2*. **b**, From the experiment in a, Na^+^ currents calculared from the time-derivative of the Na^+^ transient fits. **c**, Top, percentual change of the Na^+^ transient maximum after addition of attx (N = 8 cells) in proximal regions (mean ± SD = -1.9 ± 11.6), and in distal regions (mean ± SD = 16.8 ± 24.8); the values and statistics of the equivalent experiments with hwtx (Fig. 1) are reported for comparison. Middle, percentual change of the fast component of the Na^+^ current (I_fast_) after addition of attx in proximal regions (mean ± SD = -17.9 ± 9.9), and in distal regions (mean ± SD = 10.2 ± 13.1); the values and statistics of the equivalent experiments with hwtx are reported for comparison. Bottom, percentual change of the slow component of the Na^+^ current (I_slow_) after addition of attx in proximal regions (mean ± SD = 1.4 ± 25.2), and in distal regions (mean ± SD = 31.5 ± 49.4); the values and statistics of the equivalent experiments with hwtx (Fig. 1c) are reported for comparison; “*” indicates a significant change (p<0.01, paired t-test).

**Supplementary Figure S7.**
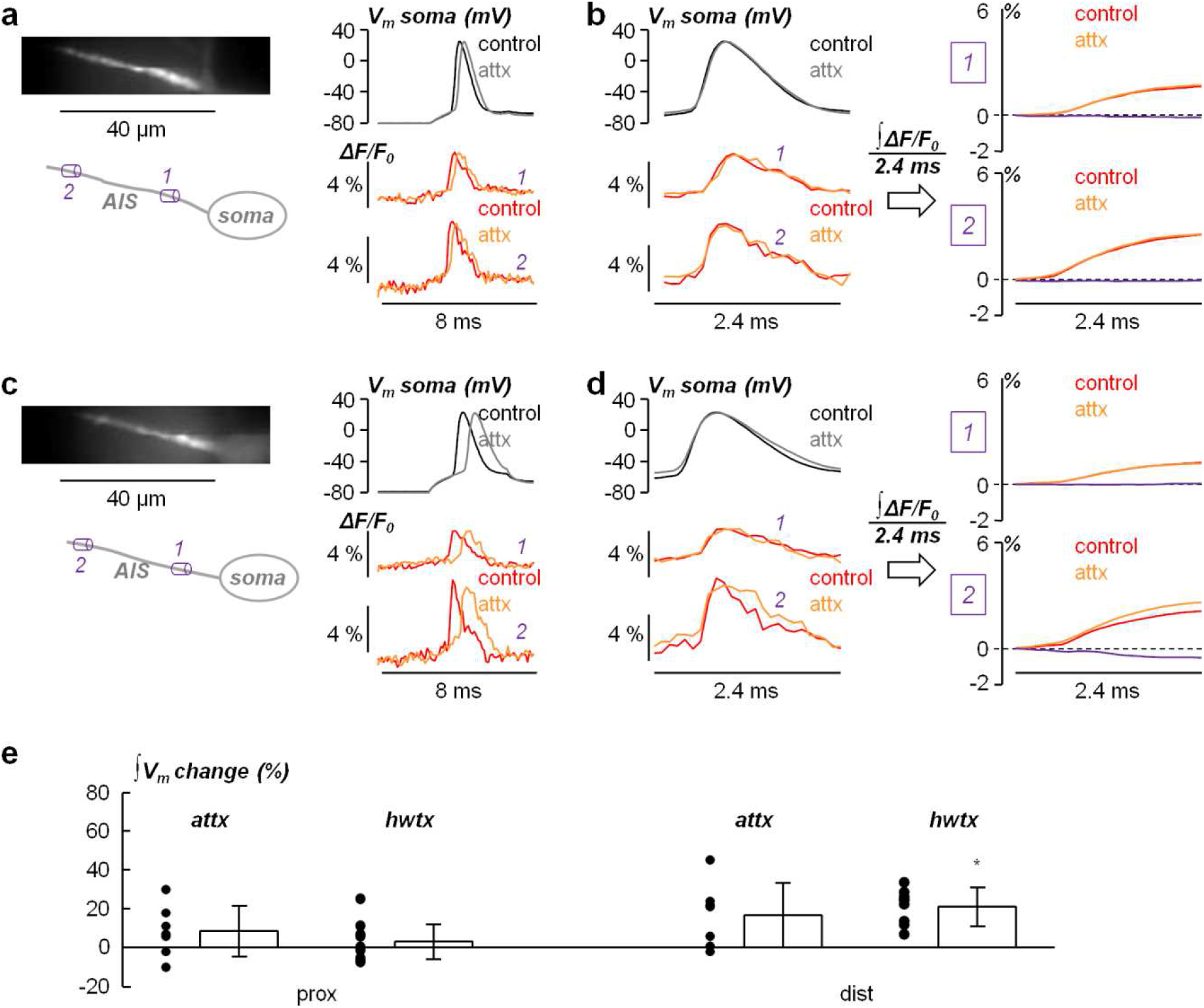
Analysis of the effect of attx on the AP waveform in the AIS. **a**, Left, fluorescence image (JPW1114) of the AIS and its reconstruction with a proximal region (*1*) and a distal (*2*) region indicated. Right, Somatic AP in control solution and after locally delivering 800 nM attx (top) and associated V_m_ transients in *1* and *2*. **b**, Left, somatic and axonal APs at a different time scale (2.4 ms duration). Right, quantification of the AP waveform shape by calculation of V_m_ integral (∫V_m_) over the 2.4 ms time window comprising the AP signal. In this cell, the local delivery of attx does not change the waveform of the axonal AP. **c** and **d**, same as a and b in another cell where the local delivery of attx widens the waveform of the axonal AP at the distal site. **e**, Percentual change of the ∫V_m_ signal maximum after delivery of attx (N=7 cells) in proximal regions (mean ± SD = 8.6 ± 13.1), and in distal regions (mean ± SD = 16.7±16.5); the values and statistics of the equivalent experiments with hwtx (Fig. 2c) are reported for comparison; “*” indicates a significant change (p<0.01, paired t-test).

**Supplementary Figure S8.**
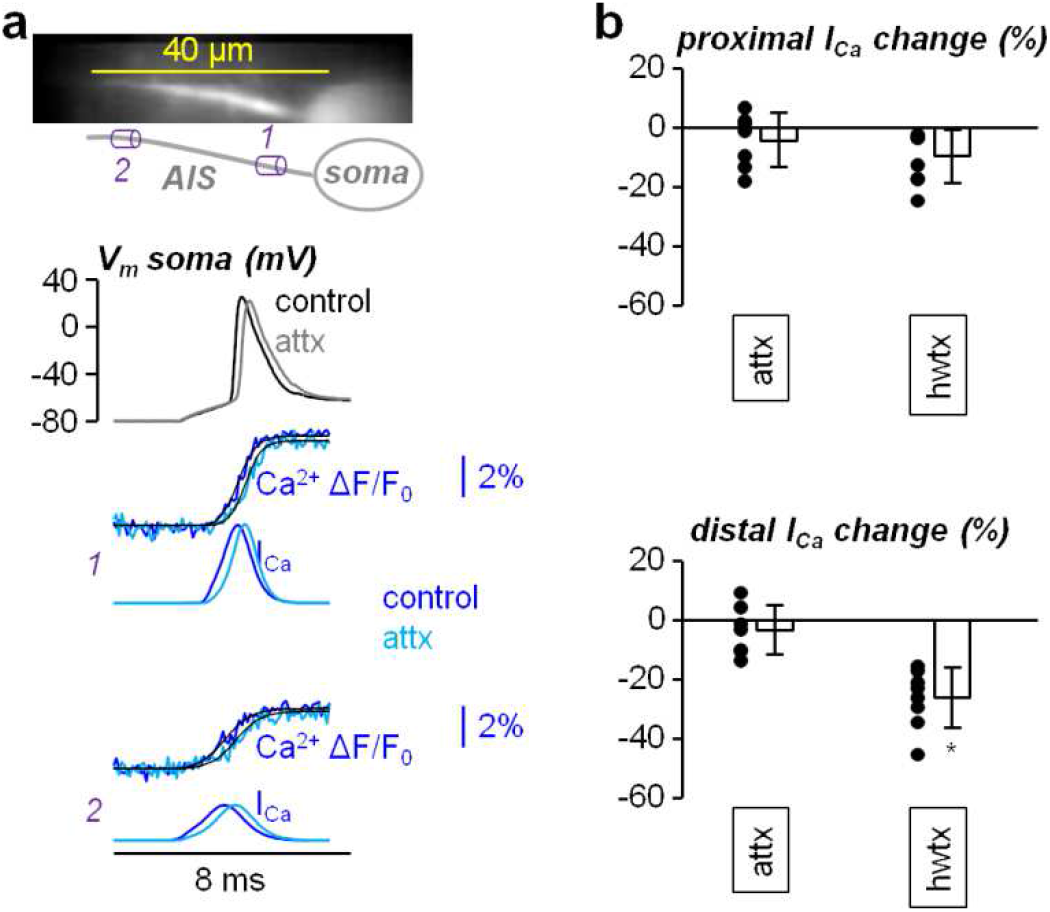
Analysis of the effect of attx on the Ca^2+^ influx in the AIS. **a**, Top, Ca^2+^ fluorescence image (OG5N) of the AIS and its reconstruction with a proximal region (*1*) and a distal (*2*) region indicated. Bottom, somatic AP and associated Ca^2+^ transients and currents in *1* and *2* in control solution and after locally delivering 800 nM attx. **b**, Top, single values and percentage change of the proximal Ca^2+^ current peak after addition of attx (N = 7 cells, mean ± SD = -4.5 ± 9.2). Right, in the same cells, single values and percentage change of the distal Ca^2+^ current peak after addition of attx (mean ± SD = -3.6 ± 8.3); the values and statistics of the equivalent experiments with hwtx (Fig. 3e) are reported for comparison; “*” indicates a significant change (p<0.01, paired t-test).

## Notes

### Competing Interest Statement

The authors have declared no competing interest.

